# Environmental impact of integrating decentralized urine treatment in the urban wastewater management system: A comparative life cycle assessment

**DOI:** 10.1101/2025.02.02.636102

**Authors:** Hanson Appiah-Twum, Tim Van Winckel, Julia Santolin, Jolien De Paepe, Stefanie Hellweg, Tove A. Larsen, Kai M. Udert, Siegfried E. Vlaeminck, Marc Spiller

## Abstract

As municipal wastewater treatment regulations become more stringent, integrating source-separated urine treatment into centralized urban wastewater management offers a ‘hybrid’ solution. However, it is not clear how the environmental impacts of such hybrid systems compare to highly efficient centralized wastewater treatment plants (WWTPs) with low N_2_O emissions and electricity use. In this study, a consequential life cycle assessment was used to compare the environmental impact of three urine hybrid wastewater treatment systems – which combine decentralized urine treatment with a highly efficient central WWTP – to a centralized WWTP treating mixed wastewater (baseline). The studied urine treatment systems include partial nitrification & distillation, struvite precipitation & stripping/scrubbing, and partial nitritation/anammox. Additionally, the contribution of urine alkalinization to the overall impact was quantified. The results show that at least one hybrid scenario showed a lower environmental impact in 8 out of the 10 assessed impact categories. Global warming potential and marine eutrophication were found to be higher than the baseline. Additionally, it was identified that urine alkalinization increased the environmental impact of the treatment system in 7 out of the 10 impact categories. A Pareto frontier analysis was developed to guide decision makers on where hybrid solutions could be used as a strategy to reduce global warming impacts of conventional WWTPs. It was realized that using N₂O emission factors of 75 WWTPs, 87% of centralized WWTPs had lower CO₂ emissions compared to partial nitrification & distillation, and 91% compared to partial nitritation/anammox hybrid solutions. However, at energy demands of 1 kWh/PE and 2 kWh/PE, both hybrid solutions showed lower emissions than all the studied WWTPs. The study highlights the potential of hybrid wastewater treatment solutions to address specific environmental challenges in wastewater management and a strategy to reduce global warming impacts in WWTPs with high N₂O emissions and electricity use.

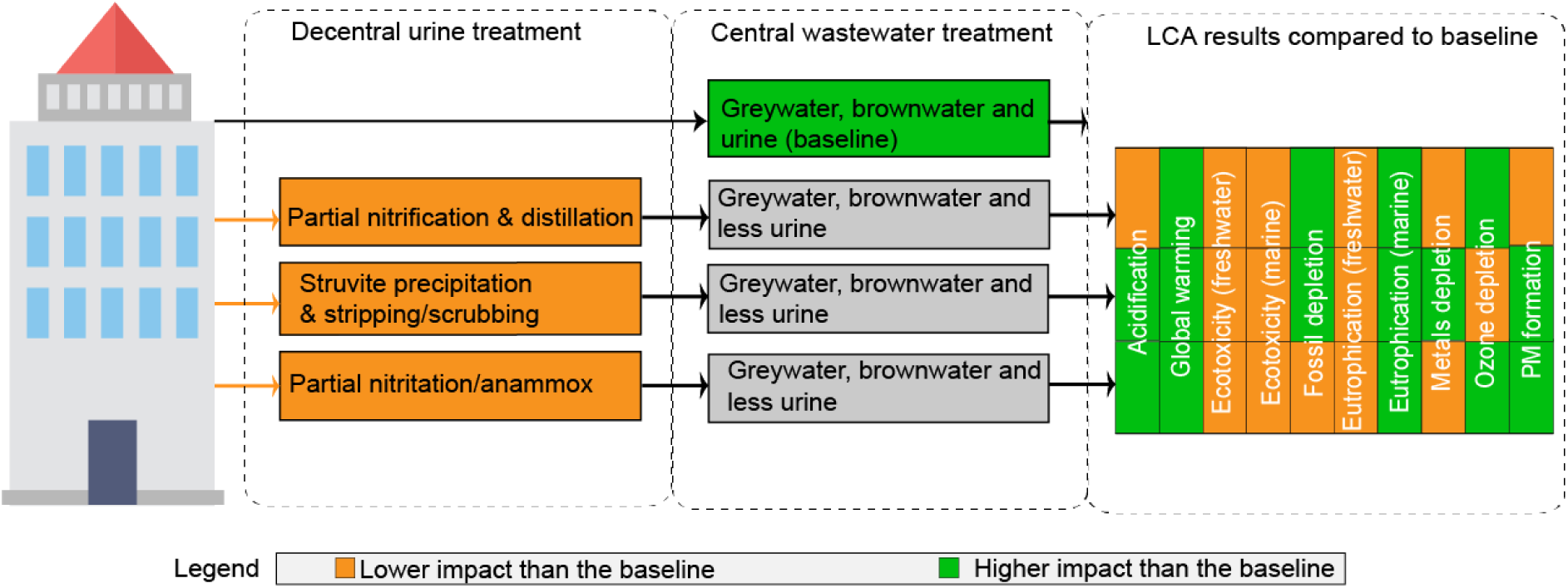
Graphical abstract.

## 1 Introduction

In the 21^st^ century, the focus of wastewater treatment has been shifting towards resource recovery, energy neutrality and climate impact reduction (Soares, 2020). For example, the EU’s urban wastewater treatment directive, urges all its member states to attain (1) a stricter nutrient removal (90% phosphorus (P) and 85% nitrogen (N)), (2) energy neutrality by 2040, (3) encourage resource recovery and (4) introduce the concept of quaternary treatment to remove 80% of indicator organic pollutants (European Commission, 2022). These requirements pose challenges for current urban wastewater systems due to the combined and diluted nature of wastewater managed by conventional wastewater treatment plants (WWTPs).

One innovation in urban wastewater management is the decentralized collection and treatment of source-separated urine (Larsen et al., 2009). Source-separation of urine presents an attractive approach, because urine constitutes less than 1% of the total volume of domestic sewage produced, but it contains on average 80% N, 50% P (Larsen & Gujer, 1996), and 64% micropollutants (Lienert et al., 2007). Therefore, it is suggested that treatment of source-separated urine can efficiently remove wastewater constituents or recover them.

Urine source-separation technologies are frequently combined with centralized black and grey water treatment in the so called “hybrid system” to achieve a reduced nutrient load to the centralized WWTP, leading to higher capacity and potentially better effluent quality (Jimenez et al., 2015). Previous research shows that these hybrid systems outperform conventional centralized WWTPs in terms of process and environmental performance through a comparative life cycle assessment (LCA). The technologies investigated in previous studies include urine storage for spontaneous fermentation and land application (Spångberg et al., 2014), struvite precipitation (Bisinella De Faria et al., 2015; Ishii & Boyer, 2015), and transmembrane chemisorption (Besson et al., 2021; Igos et al., 2017). While the majority of these studies focused on transporting separated urine to a centralized WWTP for treatment, only a few have explored the combination of decentralized urine treatment with centralized treatment of the remaining wastewater (blackwater and greywater).

Technologies such as ammonia (NH_3_) stripping in combination with struvite precipitation (Antonini et al., 2011) or the production of an ammonium nitrate (NH_4_NO_3_) through partial nitrification (Fumasoli et al., 2016) have been developed for N and P recovery. Partial nitrification followed by distillation of urine is or will be applied in several pilot installations in Europe (European Space Agency, 2023). The NH_4_NO_3_ fertilizer produced in this process has been approved as a fertilizer for edible plant production (VunaNexus, 2020). Additionally, the autotrophic partial nitritation/anammox process has been explored for removing N as N₂ gas, which is particularly suitable for urine since it is an energy efficient process and does not require an external carbon source (Udert et al., 2008). While processes such as partial nitrification, partial nitritation/anammox and NH_3_ stripping combined with struvite precipitation show promise for future applications, their environmental performance in hybrid systems, compared to conventional WWTPs, have not yet been investigated

Previous urine source-separation studies indicate that the characteristics of the centralized WWTP where N and P are treated are crucial for the outcomes of environmental assessments. In most studies, urine treatment systems are compared to WWTPs with relatively high electricity consumption and nitrous oxide (N_2_O) emissions, although these values can be highly variable between WWTPs (De Haas & Andrews, 2022; Vaccari et al., 2018). Ishii & Boyer (2015) used a central WWTP electricity demand of 1.4 kWh/m^3^ and Besson et al. (2021) applied a the IPCC N_2_O emission factor (EF) of 1.6% of influent total nitrogen (TN). Given the significance of these parameters, it is essential not only to compare innovative urine treatment technologies with highly efficient WWTPs, but also to determine the N₂O emissions at which urine source-separation results in a lower environmental impact. This is relevant because not all of the > 20,000 WWTPs in Europe (European Environmental Agency, 2023), and many more worldwide, will transition to a modern WWTP with low energy demand and N_2_O emissions.

Another knowledge gap is that studies of urine source-separation do not investigate the environmental impact of urine storage and associated NH_3_ emissions. During storage urea hydrolysis leads to a pH increase which causes NH₃ to volatilize (Randall et al., 2016). To prevent this, urine can be pre-treated during storage by inhibiting ureolysis (Udert et al., 2003). Inhibition of urea hydrolysis can be achieved through alkalinization, either by base addition (Randall et al., 2016) or electrochemical in-situ base production (De Paepe et al., 2020).

To fulfil these knowledge gaps, this study aims: (i) to determine the environmental impacts of different urine treatment systems and urine alkalinization technologies and (ii) to evaluate the effect of changes in emission at the centralised WWTP on the conclusions about environmental performance of urine source-separation. The urine treatment technologies studied are partial nitrification & distillation, struvite precipitation & stripping/scrubbing and partial nitritation/anammox. The urine pre-treatment scenarios investigated are urine alkalinization through calcium hydroxide Ca(OH)_2_ addition and electrochemical systems as well as a scenario without alkalinization (spontaneous urine fermentation). The assessment uses a consequential LCA incorporating operational processes and infrastructure data (including treatment systems and building pipes for urine transport), design of urban neighborhoods, and process modelling using real plant data.

## 2 Materials and methods

### 2.1 Goal and scope of the study

This study evaluates the environmental impact of hybrid wastewater treatment systems, which combine decentralized urine treatment with a highly efficient centralized WWTP in a Swiss city with a population equivalent (PE) of 189,000. The comparison is made against the environmental performance of the same centralized WWTP treating mixed wastewater. A consequential LCA was conducted using a functional unit of 1 PE of wastewater treated per day, ensuring compliance with the Swiss effluent discharge limits: 45 g/m³ chemical oxygen demand (COD), 0.8 g/m³ total P, 60% total N removal, 2 g/m³ ammonium (NH_4_^+^), 0.3 g/m³ nitrite (NO_2_^−^), and 20 g/m³ total suspended solids. 1 PE/day based on nutrients load in this study is defined as 12.9 g N, 1.6 g P and 136 g COD. The system boundary includes urine collection, treatment, transportation of produced fertilizer to the field and centralized wastewater treatment (Figure 1).

**Figure 1.**
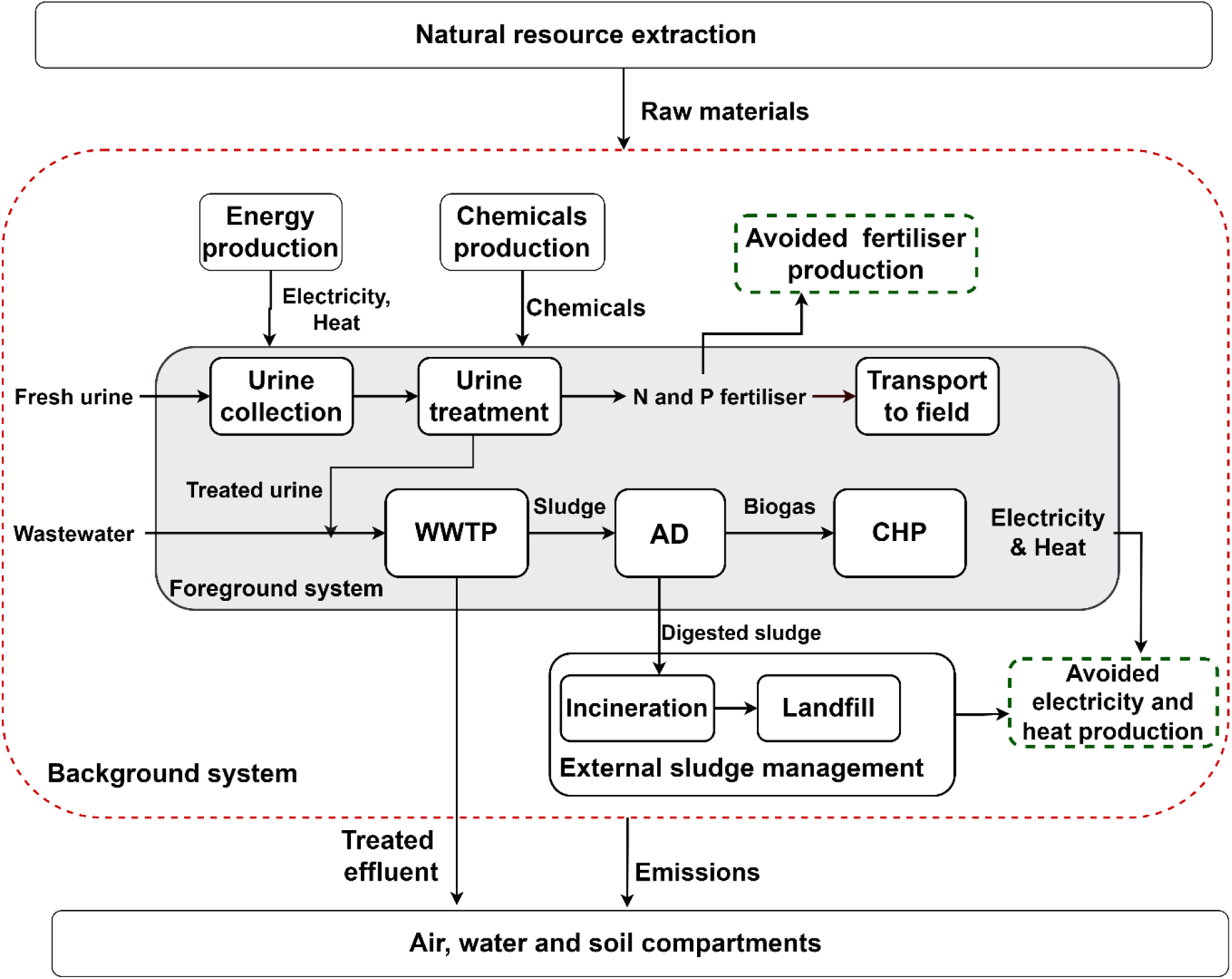
System boundary for the LCA. The large, grey-shaded box is the foreground which is inside the larger unshaded background system with a broken red boundary. WWTP: wastewater treatment plant, AD: anaerobic digester, CHP: combined heat and power reactor.

In the urine treatment scenarios, pipes for urine transportation and hardware installations for urine treatment were considered. As urine makes up less than 1% of the total hydraulic load of domestic wastewater, no changes to sewer piping or (hydraulic) dimensioning of the centralized WWTP were assumed. Therefore, infrastructure for wastewater transport and treatment at the central WWTP was excluded from the assessment, as its impact would be the same across all scenarios. Following the consequential LCA approach, system expansion was used to account for by-products from urine and wastewater treatment, such as P and N fertilizers or energy. The avoided products are single superphosphate for P and NH_4_NO_3_ for N as these are the marginal suppliers of P and N fertilizers in Europe.

### 2.2 Life cycle inventory and scenarios description

This section summarizes the inventory for the assessment. Detailed information can be found in the supplementary material.

#### 2.2.1 Studied scenarios

The baseline scenario constitutes a centralized WWTP treating mixed wastewater (without urine source-separation). This treatment is compared to three hybrid wastewater treatment systems (Table 1). The hybrid scenarios—partial nitrification & distillation, struvite precipitation & stripping/scrubbing, and partial nitritation/anammox—combine decentralized urine treatment with centralized treatment of the remaining wastewater.

**Table 1.**
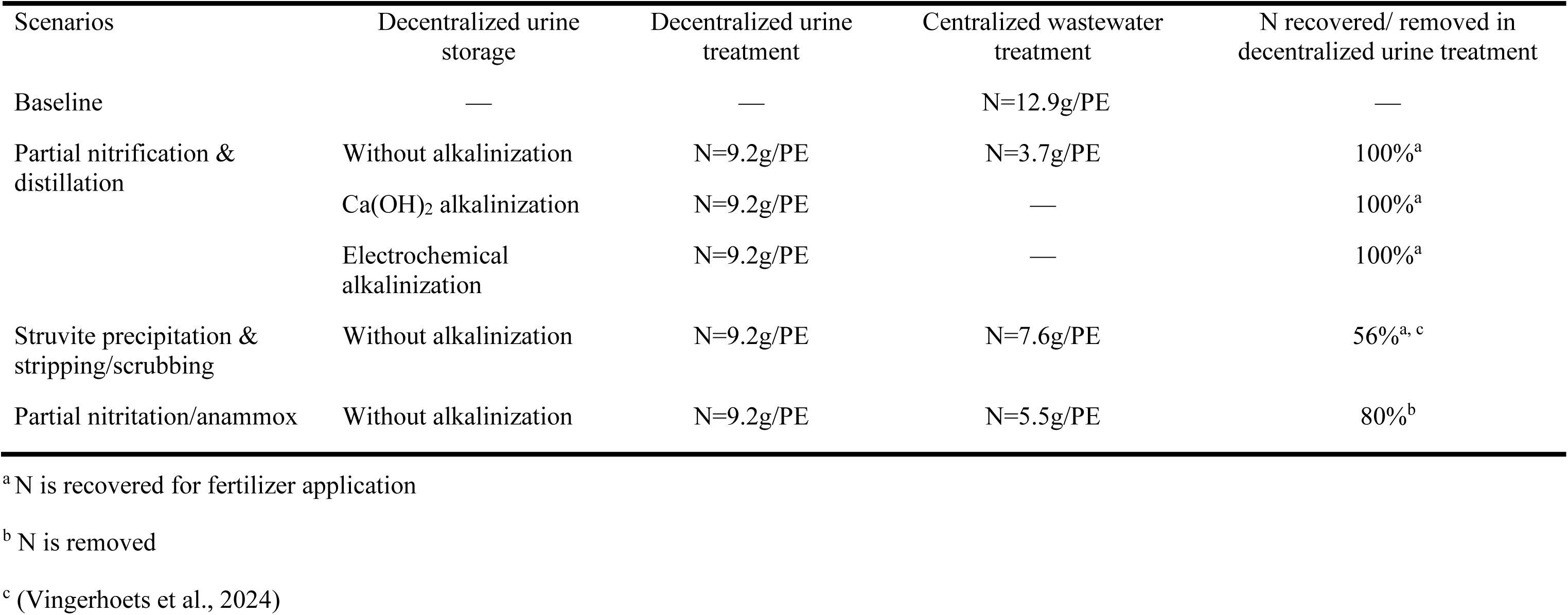
Overview of wastewater treatment scenarios, highlighting the treatment models and influent N load for decentralized urine treatment and centralized WWTPs. Nitrogen removal and recovery efficiencies for each urine treatment scenario are also presented.

An 80% urine source-separation efficiency was assumed. All urine treatment technologies received influent with the loads as specified in Table S1 (Larsen & Gujer, 1996; Udert et al., 2006). The remaining 20% urine is not treated on site but transferred through sewers to a central WWTP with the greywater, brown water and effluent of the urine treatment. The scenarios are modeled in an urban setting with a population of 189000 (the PE of the centralized WWTP) and characteristics similar to Zurich, Switzerland (Figure S1 and S2). The city is assumed to have seven-story residential buildings, each housing 400 people (PE), with an average of 2.2 PE per dwelling. Urine is collected and treated in clusters of 1200 PE, with treatment located in the basement of one building and connected to two others via 50m urine drainage pipes. The scenarios assume the entire population of the city uses source-separating toilets with separate pipe for urine transport, leading to 158 decentralized systems connected to the central WWTP in the hybrid scenarios.

The urine treatment technologies have different N removal efficiencies which impact the influent N load to the centralized WWTP (Table 1). For the urine alkalinization, three scenarios were studied for the partial nitrification & distillation decentralized urine treatment only; (i) scenario without alkalinization (ii) Ca(OH)_2_ dosage and (iii) electrochemical alkalinization. Downstream treatment of the rest of the wastewater in the centralized WWTP was excluded from the alkalinization scenarios, as it was assumed that effluent characteristics after urine treatment would be same across all scenarios.

#### 2.2.2 Centralized WWTP (Baseline)

A 365-day average of primary data from an advanced central WWTP located in Switzerland has been used in this study (Figure 2A). The biological treatment of this plant is of an A2O (anaerobic, anoxic, oxic) type, with biological N and P removal. P removal is supplemented by iron chloride (FeCl_3_) addition. Powdered activated carbon (PAC) is added for micropollutant removal and later settled and removed with the sludge. The remaining effluent is sent to a sand filtration step.

**Figure 2.**
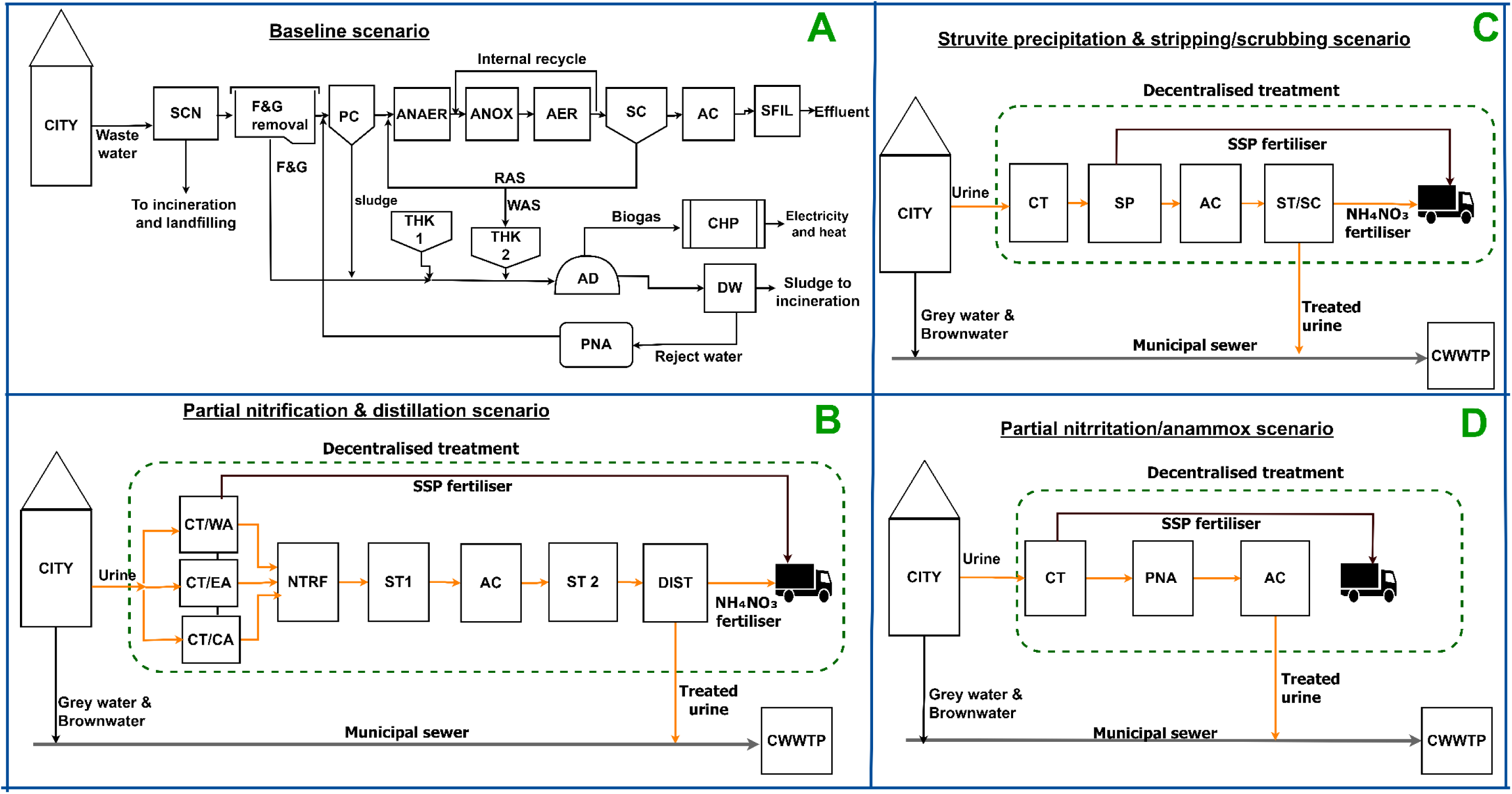
Schematic diagram of the studied scenarios. CWWTP: central wastewater treatment plant, SCN: screenings, ANAER: anaerobic tank, ANOX: anoxic tank, AER: aerobic tank, F&G: fats and grease, PC: primary clarifier, SC: secondary clarifier, AC: activated carbon reactor, SFIL: sand filtration, THK, thickener, PNA: partial nitritation/anammox, RAS: return activated sludge, WAS: waste activated sludge, DW, dewatering, AD: anaerobic digestor, CT: collection tank, NTRF: nitrification, ST: storage tank, WA: without alkalinization, EA: electrochemical alkalinization, CA: Ca(OH)_2_ alkalinization, DIST: distiller, SP: struvite precipitation, ST/SC: stripping/scrubbing, SSP: single super phosphate, NH_4_NO_3_: ammonium nitrate fertilizer.

Sludge from the clarifiers, along with fats and grease from the grease removal unit, undergo anaerobic digestion. The resulting biogas is used for electricity and heat generation in a combined heat and power (CHP) plant. All produced heat is consumed within the facility, while surplus electricity is exported to the grid, making the WWTP a net energy-positive facility (–0.001 kWh/PE/day). The digested sludge is thickened, and the liquid fraction is treated in a partial nitritation/anammox unit for N removal. The thickened digested sludge is incinerated and landfilled.

The life cycle inventory (LCI) (Table S5) generated was based on the primary data received from the WWTP operators. The total N₂O EF of the WWTP is 0.4% of influent N. This EF is a combination of 0.3% of influent N from the activated sludge (primary data) and 1.8% of N load in reject water from treatment in a partial nitritation/anammox (Dieziger et al., 2023). The other emissions include methane (CH_4_) from the activated sludge (0.1% influent COD) and sludge management (0.9% influent COD), and indirect N_2_O emissions from sludge incineration (3.8% N in sludge) (Gruber, 2021). All direct CO_2_ emissions were considered biogenic and were excluded from the assessment.

#### 2.2.3 Urine collection and treatment infrastructure

The in-building piping infrastructure for urine transport was modelled in accordance with the International Plumbing Code (IPC, 2024) (Section 3, Tables S2 and S3 of Supplementary Material). PVC pipe material was assumed for the pipes. For underground residential sewer lines connecting the three buildings to the treatment facility, the Ecoinvent process “residential sewer grid construction, 0.087 km, CH” was used. Infrastructure data for the treatment systems were assumed to have a 20-year lifespan for all unit process hardware (Table S4).

#### 2.2.4 Decentralized urine storage and alkalinization

Source-separated urine is stored in collection tanks (one per three buildings) before treatment, with an assumed dilution factor of 2.6 as a result of flushing (Faust et al., 2022). In scenarios without urine alkalinization, urine is stored under spontaneous fermentation conditions, leading to a pH rise to over 9 and potential NH₃ emissions, which are assumed to be minimized through the use of a one-way air valve (as implemented in the pilot in Switzerland). An NH₃ EF of 0.5% of influent N in the storage tank was assumed.

In urine alkalinization scenarios (Table S6), NH₃ emissions are assumed to be negligible. In the electrochemical alkalinization scenario (De Paepe et al., 2020), the urine pH is raised to approximately 12, inhibiting urea hydrolysis, with an energy requirement of 14 Wh/L and achieving 70% P removal as calcium phosphate through precipitation. For Ca(OH)₂ alkalinization (Randall et al., 2016), 10g Ca(OH)_2_ is added per liter of urine and stirred electrically leading to precipitation of all phosphate as calcium phosphate.

To prevent scale formation and cross-contamination during cleaning of the source-separating toilet, a 10% citric acid solution is used. Based on experiences from Eawag, bi-weekly cleaning with 40 ml of this solution is assumed for all decentralized urine treatment scenarios, except the electrochemical alkalinization scenario whose electrolyte serves as a cleaning detergent.

#### 2.2.5 Partial nitrification & distillation scenario

The partial nitrification & distillation scenario (Figure 2B) is based on information from a pilot installation located in Dübendorf (Eawag, Switzerland) (Faust et al., 2022) (Table S6). After storage, hydrolyzed urine is partially nitrified in a suspended growth reactor, resulting in a 50/50 NH_4_-N/NO_3_-N ratio, with N₂O emissions of 1.5% of the influent N load. The solution is then treated with granular activated carbon (GAC) for micropollutant removal (569 mg GAC/L urine) (Köpping et al., 2020) before being concentrated 10-15 times in a distillation unit. The amount of GAC used is the same across all urine treatment scenarios. The final product is an NH_4_NO_3_ fertilizer with a composition of 42 g TN and 4 g phosphate measured as P_2_O_5_ in a liter of the fertilizer product that is approved for use in agriculture/horticulture in Switzerland, Lichtenstein, France and Austria (VunaNexus, 2020). The only process effluent is the distiller condensate, used for toilet flushing at Eawag but assumed to be directly discharged to the central WWTP in this study.

For the urine alkalinization pre-treatment, it was estimated that the high alkalinity of the hydrolyzed urine leads to 60% and 86% nitrification rate in the electrochemical and Ca(OH)_2_ scenarios respectively (i.e. the nitrification process is stopping when all alkalinity is consumed due to pH drop). Subsequently the electricity demand for nitrification for the electrochemical (7.1 Wh/L urine) and Ca(OH)_2_ (10.1 Wh/L urine) were estimated based on (Faust et al., 2022).

#### 2.2.6 Struvite precipitation & stripping/ scrubbing scenario

In this scenario, (Figure 2C) hydrolyzed urine enters a struvite reactor, where magnesium oxide (MgO) is added to the urine in a molar ratio of 1.5 Mg:1 P (Antonini et al., 2011). This process recovers 98% P as struvite, with no additional base required due to the suitable pH after urea hydrolysis (Table S7). In the next step, the liquid passes over a GAC unit for micropollutant removal before entering an air stripping unit. The air stripping unit (Vingerhoets et al., 2024) operates at pH 8.5-8.6 and 50°C temperature. Air with a flow rate of 0.2–0.8 m s^−1^ is blown through the liquid, transferring approximately 56% of the N into the gas phase as NH₃. This gas is then scrubbed with 60% nitric acid, forming NH_4_NO_3_ fertilizer. The combined struvite precipitation & stripping/ scrubbing resulted in an estimated 0.085 kWh/PE electricity use. The remaining liquid, containing 2% of the initial P and 44% of the N, is discharged to the centralized WWTP.

#### 2.2.7 Partial nitritation/ anammox

In this scenario (Figure 2D) about 50% of the NH_3_ in the stored urine is oxidized to nitrite and then reduced to N_2_ gas using anammox bacteria. The process is established for side stream wastewater treatment, but has been applied to urine using a sequencing batch reactor (Udert et al., 2008). Energy demand was estimated based on (Faust et al., 2022). The estimated electricity including pumping and process control was 0.025 kWh/PE. An N_2_O EF of 1.8% of influent was used (Dieziger et al., 2023). The treated urine (containing 20% N and 80% P of the source-separated urine) is then discharged via a GAC to the WWTP (see Table S8 for the LCI).

#### 2.2.8 Centralized wastewater treatment modelling approach

Using primary data from the WWTP, a steady-state model of the centralized WWTP was developed in SUMO wastewater modeling software (Dynamita, version 22.0.0) (Table S9). Unknown parameters, such as aeration efficiency, were calibrated against these primary data. This calibrated model was then used to simulate changes in influent composition due to urine source-separation. To meet the Swiss effluent discharge limits, adjustments were made to the aeration model, blower efficiency, air input, and chemical dosing. The simulation results served as proxy data for the LCA (Table S5).

In the PAC reactor for micropollutant removal, electricity, chemical and material consumption (activated carbon, FeCl₃, and polymer) were reduced by 49% based on findings by Lienert et al. (2007), who indicated that 64% of micropollutants in wastewater originate from urine (i.e. 100-64=36+(64*(1-0.8)=48.8%)) (Lienert et al., 2007). The sludge line and electricity demand for other unit processes were not recalibrated, as no significant changes were expected. Similarly, biogas, electricity, and heat production were assumed to remain unchanged after urine source-separation.

### 2.3 Life cycle impact assessment

Activity Browser software version 2.6.9 and the Ecoinvent consequential database 3.9.1 were used for the LCA modelling. Environmental impacts were assessed using the ReCiPe 2016 midpoint (H) method (Huijbregts et al., 2017). Ten impact categories were studied based on their relevance to the goal and scope of the study. These include acidification, ecotoxicity (freshwater), ecotoxicity (marine), eutrophication (freshwater), eutrophication (marine), fossil depletion, global warming, minerals depletion, ozone depletion and particulate matter formation.

### 2.4 Sensitivity analysis

A sensitivity analysis was conducted on direct N_2_O emissions, electricity use and different electricity mix as these were found to be the most influential parameters on global warming impact.

A Pareto frontier analysis was conducted to determine the direct N_2_O emissions at which the baseline and hybrid scenarios have equal global warming impact. First, the impact of the baseline centralized WWTP, excluding direct N₂O emissions (*a*) was calculated. For the hybrid scenarios, the impacts of the centralized WWTP (*b*) and decentralized urine treatment (*c*) were determined also excluding direct N₂O emissions.

The total global warming impact of the centralized WWTP for both baseline and hybrid scenarios were then evaluated at various direct N₂O EFs using Equation (1) and (2) respectively. Similarly, the global warming impact of decentralized urine treatment was assessed at different N₂O EFs using Equation (3).

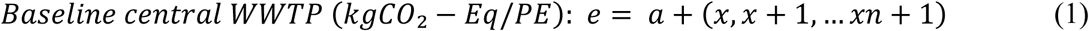

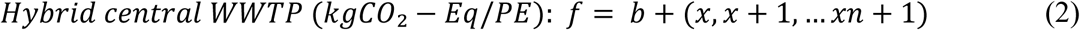

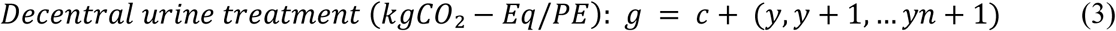

Where x and y are the kgCO_2_-Eq from direct N_2_O emissions (% influent N) of centralized WWTP and decentralized urine treatment respectively.

Next, a matrix addition of *f* and *g* was performed to determine the total global warming impact for hybrid scenarios across different N₂O EFs. A cross-check was then conducted to identify the N₂O EFs at which the global warming impacts of the baseline (*e*) and hybrid scenarios (*f + g*) were equivalent. Finally, the N₂O EFs at which the baseline and hybrid scenarios achieved equivalent impacts were plotted.

This analysis was done for different electricity use of the central WWTP; –0.001 kWh/PE electricity use from this study and assumed values ranging from 0.5 to 2 kWh/PE. Struvite precipitation & stripping/scrubbing was excluded from this analysis as it does not directly emit N₂O in the urine treatment process.

Furthermore, the marginal electricity mix in three countries (Hungary, Croatia and India) were considered to evaluate how sensitive the results are to different electricity sources. The electricity mix of these countries was selected based on the carbon intensity of their electricity generation, as projected in the Ecoinvent consequential database 3.9.1. Hungary (0.016 kgCO_2_-Eq/kWh) and Croatia (0.725 kgCO_2_-Eq/kWh) were selected due to their lowest and highest potential impacts on global warming in Europe, respectively. India (1.248 kgCO_2_-Eq/kWh) was selected based on its highest carbon intensity in the database.

## 3 Results

### 3.1 Life cycle inventory analysis

#### 3.1.1 Analysis of the baseline and hybrid scenarios’ process performance

The baseline central WWTP has a gross electricity demand of 0.059 kWh/PE/day (0.18 kWh/m^3^) with aeration for nutrient removal in the activated sludge process accounting for approximately 33% of this amount (Figure 3A). However, its net electricity consumption is –0.001 kWh/PE/day, meaning it generates more electricity than it consumes.

**Figure 3.**
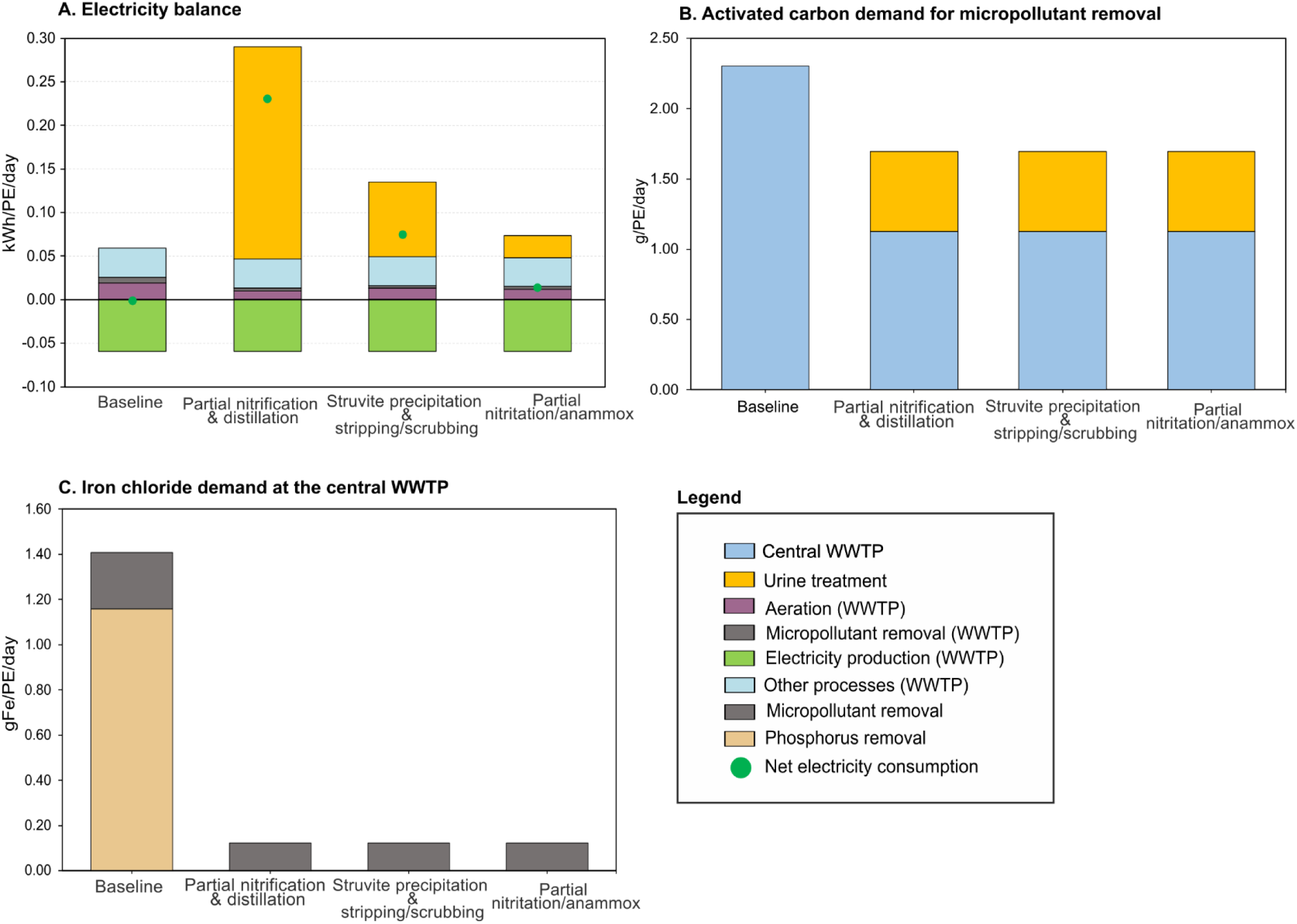
Inventory results of the baseline and hybrid scenarios. A: electricity balance, B: activated carbon demand and C: iron chloride demand at the central WWTP. The green dot represents the net electricity use due to electricity production at the central WWTP

The treated effluent complies with the discharge limits outlined in Section 2.1. 2.3 g Fe/PE/day activated carbon is used for micropollutant removal in the central WWTP (Figure 3B). FeCl₃ is applied for chemical precipitation at a rate of 1.2 g Fe/PE/day to supplement biological P removal and 0.25 g Fe/PE/day in the micropollutant removal reactor (Figure 3C).

All the hybrid solutions presented increased electricity consumption compared to the baseline scenario (Figure 3A). The partial nitrification & distillation (without alkalinization) scenario had the highest electricity use (0.231 kWh/PE/day), with 87% of this attributed to the distillation process in urine treatment. This was followed by struvite precipitation & stripping/scrubbing (0.075 kWh/PE/day) and partial nitritation/anammox (0.014 kWh/PE/day) scenarios. The hybrid scenarios’ electricity consumption values were 16 to 265 times higher than the baseline.

With 80% urine source-separation, the hybrid treatment scenarios decrease aeration electricity demand at the central WWTP by factors of 1.9 (partial nitrification & distillation), 1.6 (partial nitritation/anammox), and 1.5 (struvite precipitation & stripping/scrubbing). Furthermore, electricity demand for micropollutant removal in the central WWTP is reduced by half in these hybrid scenarios compared to the baseline. However, despite these energy savings at the central WWTP, the baseline scenario maintained a lower o electricity demand due to the high electricity requirements of urine treatment in the hybrid scenarios.

Results from the urine alkalinization scenarios indicate that urine alkalinization raises the electricity demand for urine treatment (Figure S4). Specifically, electricity consumption in the Ca(OH)₂ and electrochemical scenarios was 1.05 and 1.16 times higher, respectively, than in the scenario without alkalinization. This increase is attributed to the electricity required for the alkalinization process in the electrochemical scenario and the increased aeration demand in nitrification due to higher alkalinity levels.

#### 3.1.2 Influence of urine source-separation on chemical consumption

The study demonstrates that urine source-separation decreases FeCl₃ and activated carbon consumption in wastewater treatment (Figure 3). In the hybrid scenarios, urine source-separation eliminated the need for FeCl₃ for P removal to meet the effluent limits, while also reducing FeCl₃ usage for micropollutant removal by 49%. Additionally, activated carbon demand for micropollutant removal was lowered by 27% in the hybrid scenarios.

#### 3.1.3 Influence of urine source-separation on effluent quality

All scenarios complied with the effluent discharge limits outlined in Section 2.1 (Table 2). While COD and total P concentrations were similar between the baseline and hybrid scenarios, the total N concentration was lower in the hybrid scenarios by almost a factor of three compared to the baseline. An examination of the components of total N (NH_4_^+^, nitrate (NO_3_^−^), and NO_2_^−^) showed that the baseline scenario had a lower NH_4_^+^ effluent concentration (about 9 times lower) compared to the hybrid scenarios. This difference is attributed to a decreased concentration of nitrifiers in the centralized WWTP under the hybrid scenarios (Figure S5). Consequently, the hybrid scenarios had lower NO_2_^−^ and NO_3_^−^concentrations due to decreased nitrification activity.

**Table 2.** Effluent quality of the baseline and hybrid scenarios.

| Parameters | Unit | Baseline | Partial nitrification & distillation | Struvite precipitation & stripping/scrubbing | Partial nitrification/anammox |
| --- | --- | --- | --- | --- | --- |
| COD | (g/m<sup>3</sup>) | 11.5 | 11.2 | 12.0 | 11.1 |
| TN | (g/m<sup>3</sup>) | 5.8 | 2.3 | 2.1 | 2.1 |
| NH<sub>4</sub> | (g/m<sup>3</sup>) | 0.2 | 1.9 | 1.8 | 1.8 |
| NO<sub>2</sub><sup>-</sup> | (g/m<sup>3</sup>) | 0.07 | — | — | — |
| NO<sub>3</sub><sup>-</sup> | (g/m<sup>3</sup>) | 5.1 | — | — | — |
| Organic N | (g/m<sup>3</sup>) | 0.42 | 0.45 | 0.33 | 0.27 |
| TP | (g/m<sup>3</sup>) | 0.1 | 0.1 | 0.1 | 0.1 |
| TSS | (g/m<sup>3</sup>) | 1.0 | 1.0 | 1.0 | 1.0 |

### 3.2 Impact assessment results of the baseline, three hybrid and urine alkalinization scenarios

At least one hybrid scenarios had a lower impact than the baseline in 8 out of 10 categories, except for global warming and marine eutrophication (Figure 4). The baseline scenario recorded the highest impacts in three categories: freshwater ecotoxicity, marine ecotoxicity, and freshwater eutrophication.

**Figure 4.**
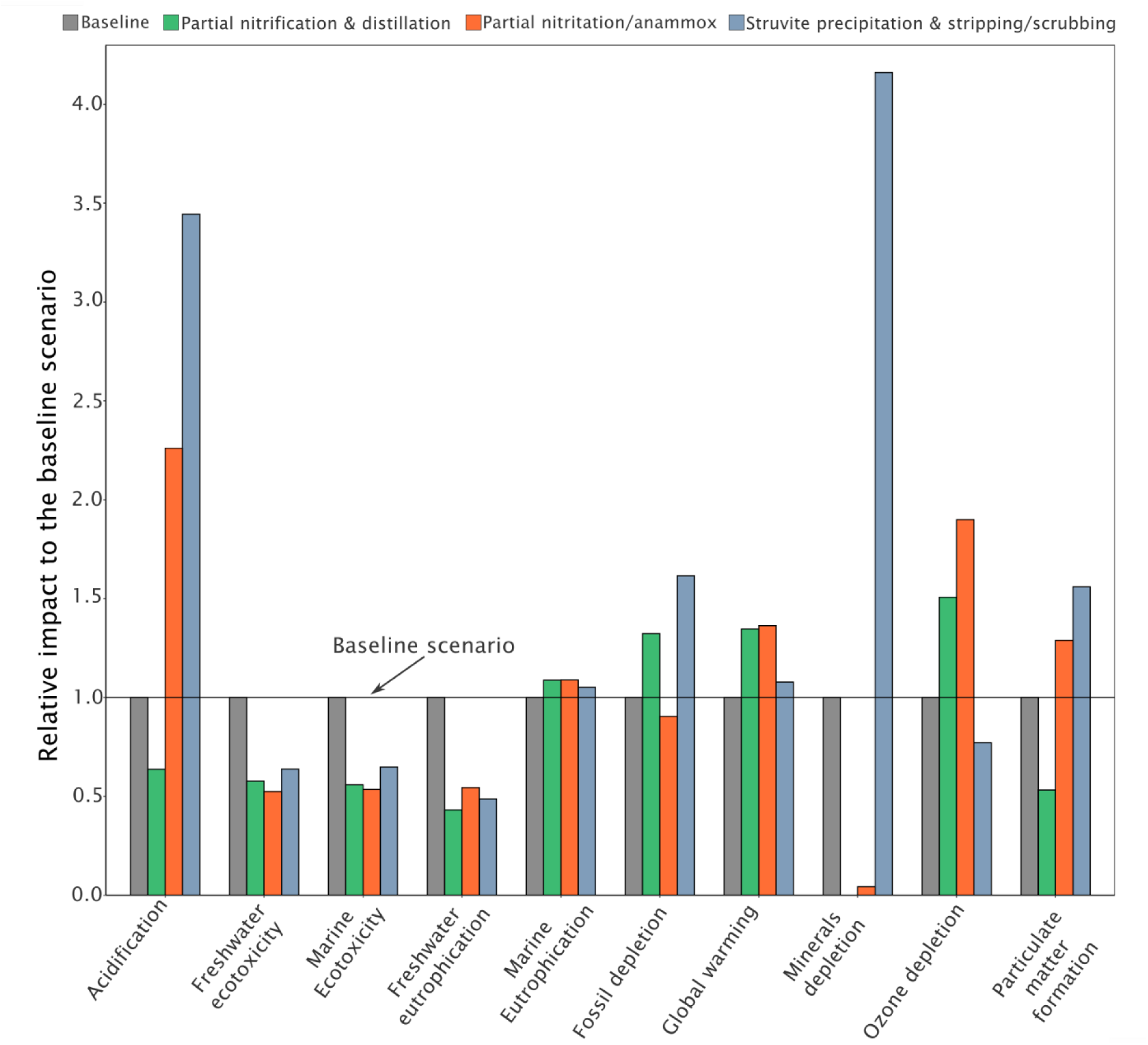
Impact assessment results of the hybrid wastewater treatment scenarios relative to the baseline. An impact above 1 indicates a higher impact than the baseline while and impact below 1 indicates a lower impact than the baseline (see Table S10 for net impacts).

The partial nitrification & distillation scenario presented the lowest impact in four categories (acidification, freshwater eutrophication, minerals depletion and particulate matter formation) and the highest in none. Partial nitritation/anammox showed the lowest impact in three categories including freshwater and marine ecotoxicity and fossil depletion. However, impacts in marine eutrophication global warming, and ozone depletion were the highest among the studied scenarios. The struvite precipitation and stripping/scrubbing scenario showed the lowest impact only in ozone depletion, but it had the highest impacts in four categories: particulate matter formation, fossil depletion, acidification, and minerals depletion, with impacts reaching 1.6 to 4 times that of the baseline.

In the urine pre-treatment scenarios, the scenario without alkalinization had the lowest environmental impact in six categories including global warming and the highest in three (acidification, freshwater eutrophication and ozone depletion) (Table S11). Ca(OH)_2_ and electrochemical alkalinization showed reduced impact in the acidification by a factor of 11 to 12 times the impact of the scenario without alkalinization.

### 3.3 Contribution analysis

#### 3.3.1 Analysis of the global warming contribution in the baseline and hybrid scenarios

The global warming impact in the baseline was 0.142 g kgCO_2_-Eq/PE/day, while in the hybrid scenarios it ranged from 0.153 to 0.193 kgCO_2_-Eq/PE/day (Figure 5).

**Figure 5.**
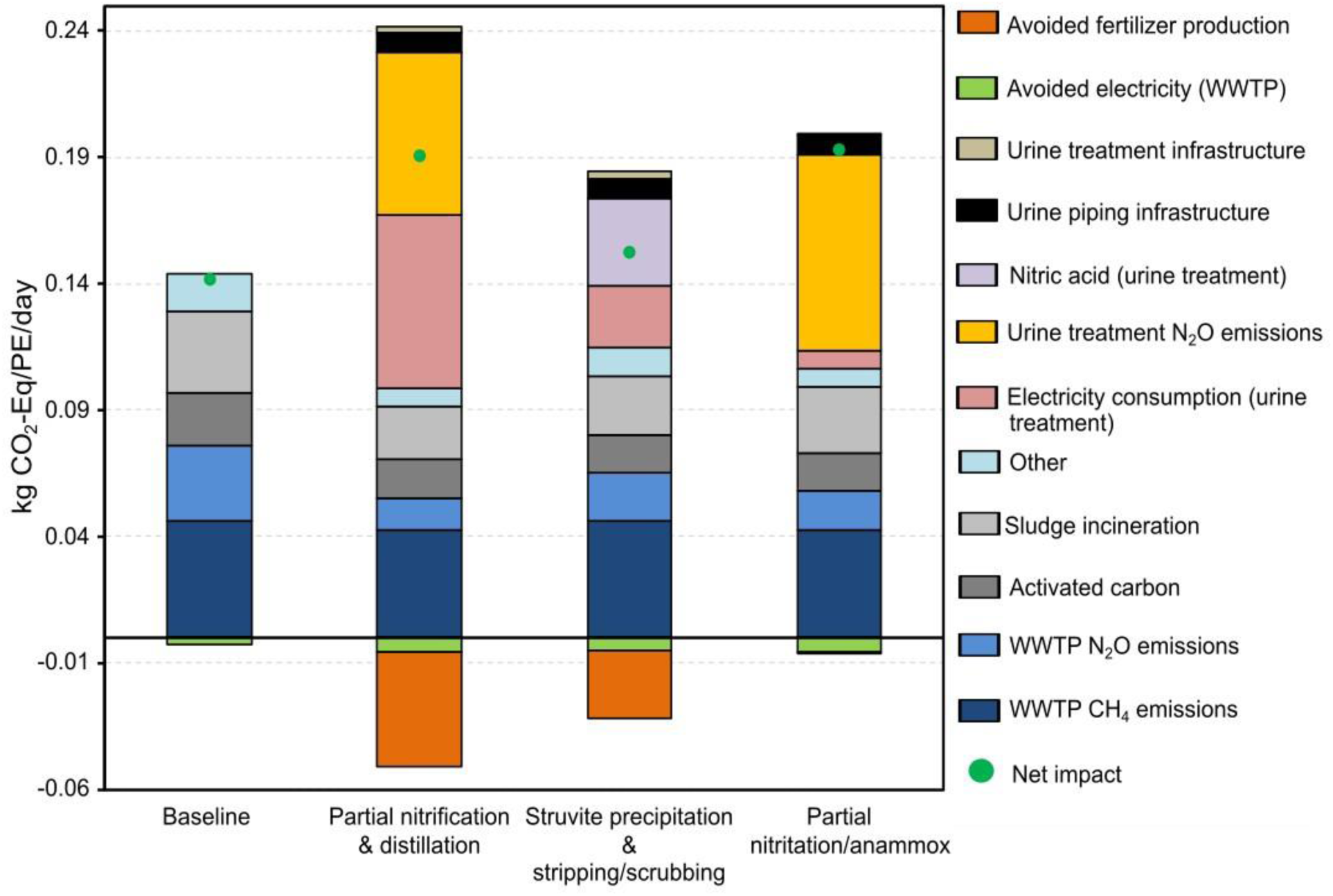
Contribution analysis of the studied scenarios’ impact on global warming. All scenarios are without alkalinization pre-treatment.

In the baseline, the impacts were dominated by direct CH_4_ emissions from the central WWTP, contributing 33% of the total impacts. Indirect N_2_O emissions from sludge incineration contributed the second highest impact (23%) followed by direct N_2_O emissions (21%) from the activated sludge and side stream treatment of reject water.

In the hybrid scenarios, a reduction in nutrient load to the central WWTP saw a reduction in the global warming impact of the central WWTP by a factor of 1.4 to 1.7 times the baseline scenario’s impact. This reduction is due to decreased N_2_O emissions and electricity and activated carbon consumption. Impact from CH₄ emissions from the central WWTP remained nearly unchanged across the hybrid scenarios, as the amount of COD entering the WWTP was similar to the baseline.

However, the overall impact of the hybrid scenarios exceeded the baseline due to impact from decentralized urine treatment. In the partial nitrification & distillation scenario, 28% of the impact was due to electricity consumption by the distillation unit and 27% from N₂O emissions. For struvite precipitation & stripping/scrubbing, 19% of the impact was from nitric acid use and 13% from electricity. In the partial nitritation/anammox scenario, N₂O emissions from urine treatment contributed 39% of its impact. A combined avoided fertilizer and electricity production reduced the global warming impact in the hybrid scenarios by 4% to 20% in the hybrid scenarios.

Overall, the impacts from infrastructure (piping and treatment system) were very low (5%) compared to the operational impacts.

#### 3.3.2 Marine eutrophication

The marine eutrophication impact in the baseline scenario was 0.131 g N-Eq/PE/day, while in the hybrid scenarios it ranged from 0.138 to 0.143 g N-Eq/PE/day (Figure 6). In the baseline scenario, the impact was dominated by NO₃⁻ (86%) and NH₄⁺ (12%) in the treated water discharged from the central WWTP. In the hybrid scenarios, 97% to 98% of the impact was due to NH₄⁺ in the discharged effluent. The increased NH₄⁺ concentrations in the hybrid scenarios’ effluent contributed to their higher overall impact compared to the baseline. Avoided impacts from fertilizer production was very low across the hybrid scenarios, ranging from 0.1% to 3.5%.

**Figure 6.**
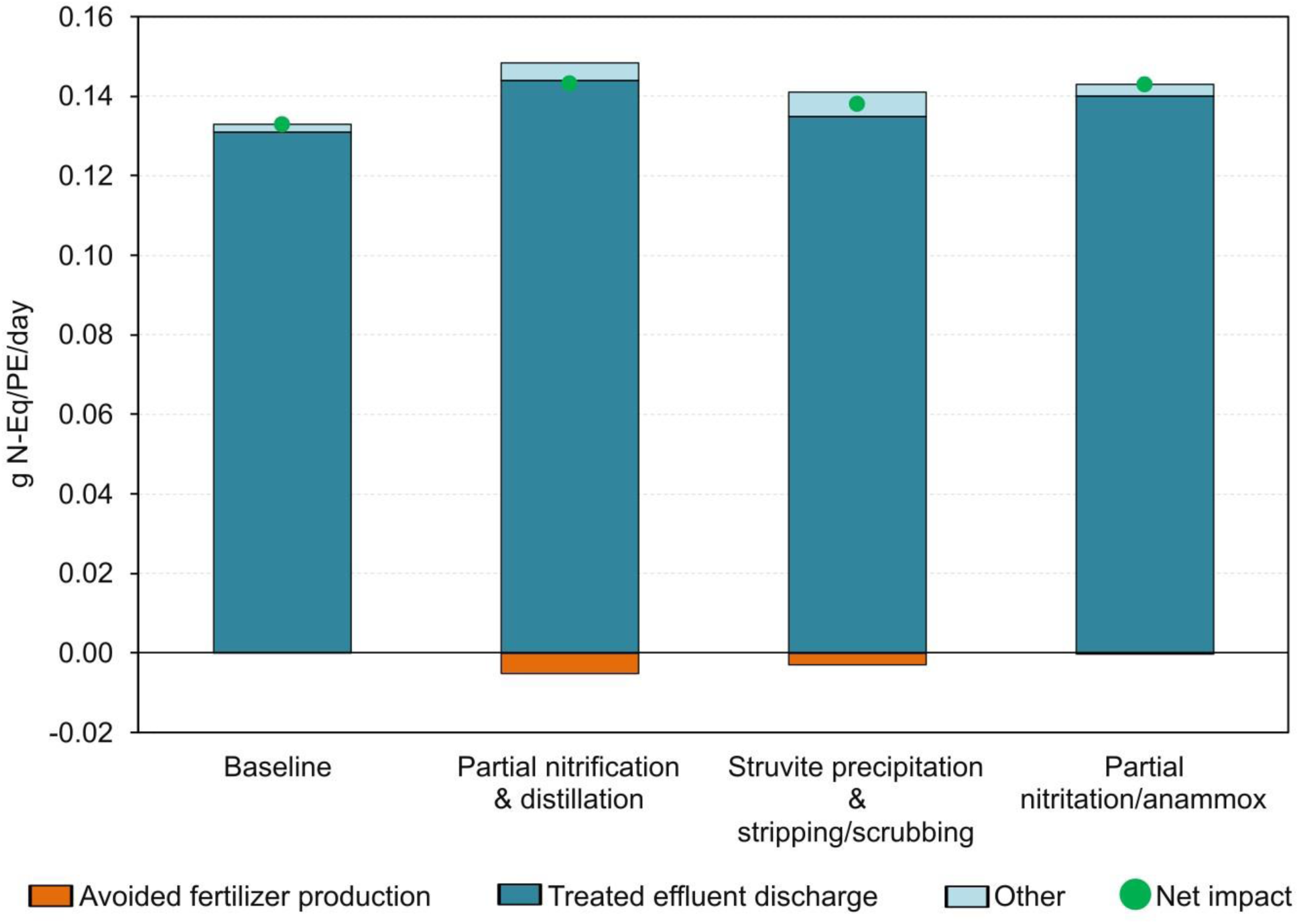
Contribution analysis of marine eutrophication impact for the baseline and hybrid scenarios.

### 3.4 Sensitivity analysis of N_2_O emissions and electricity mixes impact on global warming

#### 3.4.1 Sensitivity analysis of direct N_2_O emissions impact on global warming

The Pareto frontier analysis (Figure 7) shows the tradeoffs between direct N₂O emissions and other GHGs in the baseline and hybrid scenarios. The Pareto front represents the points where the global warming impacts of the baseline and hybrid scenarios are equal at specific N₂O emission levels.

**Figure 7.**
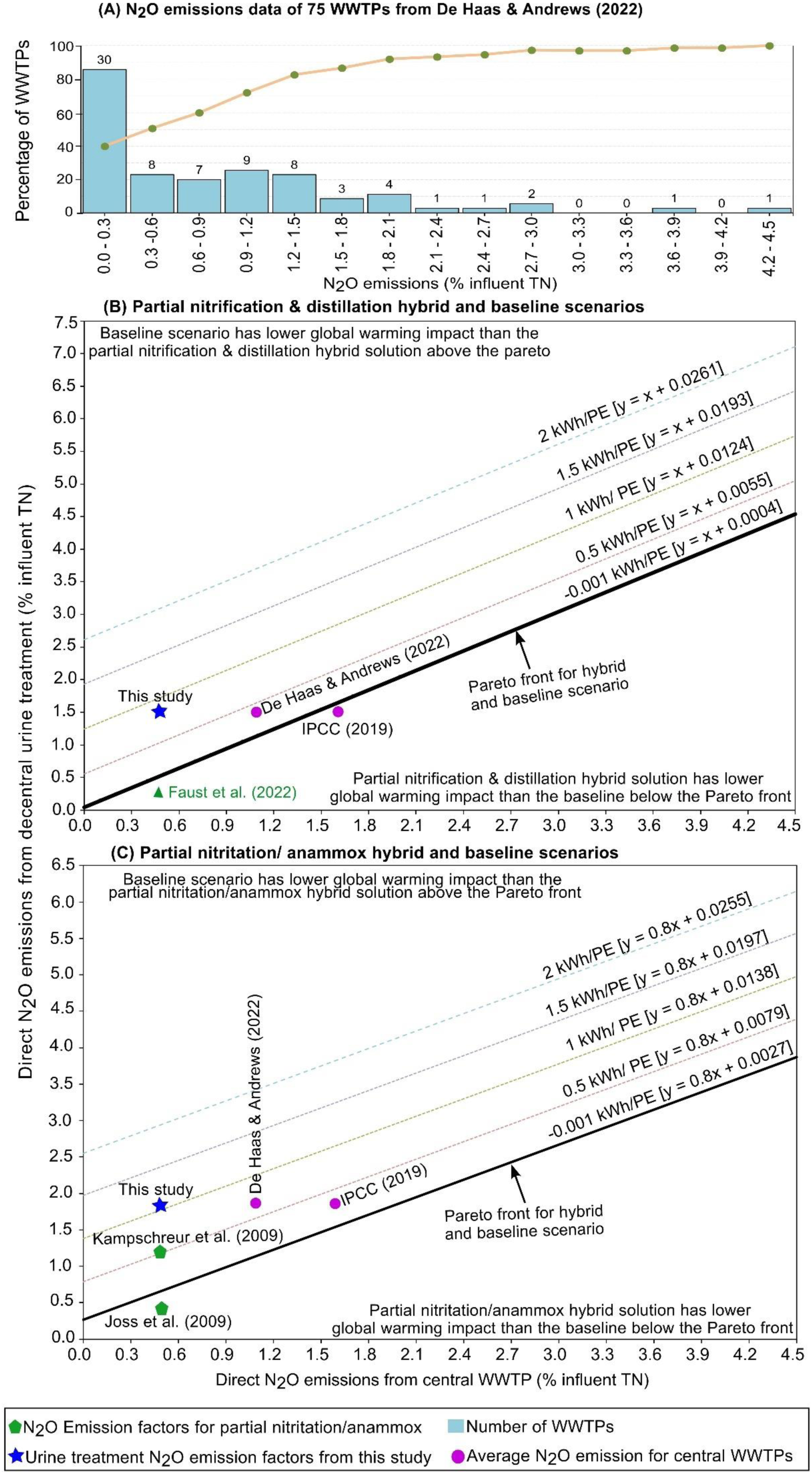
Tradeoffs between direct N₂O emissions and other GHGs in the baseline and hybrid scenarios. On the Pareto front, both the baseline and hybrid scenarios have the same impact. Above the Pareto front, the hybrid scenarios have a higher global warming impact than the baseline and vice versa. Frame A: N_2_O emissions of 75 WWTPs from De Haas & Andrews (2022); B: Pareto frontier analysis for partial nitrification & distillation hybrid and baseline scenarios and C: Pareto frontier analysis for partial nitritation/anammox hybrid and baseline scenarios.

Comparing partial nitrification & distillation hybrid and baseline scenarios shows a slope of 1 (Figure 7B). This indicates that for each unit increase in WWTP direct N₂O emissions, the urine treatment emissions must increase by an equivalent amount to maintain the same global warming impact. In contrast, the slope for partial nitritation/anammox hybrid and baseline scenarios is 0.8 (Figure 7C). This result implies that a unit increase in WWTP direct N₂O emissions requires only an 80% increase in the decentralized urine treatment emissions to achieve an equivalent impact. This difference between the two scenarios arises because partial nitrification & distillation recover all N from the separated urine, whereas partial nitritation/anammox removes 80% of the N in the source-separated urine. As a result, partial nitrification & distillation lead to a relatively lower N load in the wastewater influent to the centralized WWTP.

The results further show that N₂O EFs of about 87% of 75 WWTPs are below 1.54%, the value at which the current EF for partial nitrification and distillation (1.5%) has lower CO2-eq. emissions (Figure 7A). For the partial nitritation/anammox scenario, which assumes an EF of 1.8%, the global warming impact is higher than 91% that of WWTPs. However, reducing the N_2_O EFs for the urine treatment systems changes the outcomes. For instance, if the EF for partial nitrification & distillation is lowered to 0.26% (Faust et al., 2022), then the number of WWTPs with lower emissions reduces to 32%. Similarly, adopting an EF of 0.4% for partial nitritation/anammox (Joss et al., 2009) reduces the proportion of WWTPs with lower emissions from 91% to 28%.

These findings, however, are specific to the highly energy-efficient WWTP studied in this work. If the electricity demand of the WWTPs increases, ranging between 0.5 and 2 kWh/PE and assuming the Swiss electricity mix (Figure 8), the results change. An increase in energy demand shifts the frontier leftward and raises the y-intercept, making hybrid solutions more advantageous even at lower central WWTP N_2_O EFs. For example, a 0.5 kWh/PE increase in electricity demand raises the y-intercept by 0.70% EF for partial nitrification-distillation and by 0.60% EF for partial nitritation/anammox. This means that when WWTP electricity demand increases by 0.5 kWh/PE, the partial nitrification & distillation and partial nitritation/anammox hybrid can increase their N₂O EFs to 0.7% and 0.6%, respectively, while maintaining equivalent impacts at a specific WWTP N₂O EF. At an electricity demand of 2 kWh/PE, the y-intercept increases to values ranging between 2.55% and 2.61% for both scenarios.

**Figure 8.**
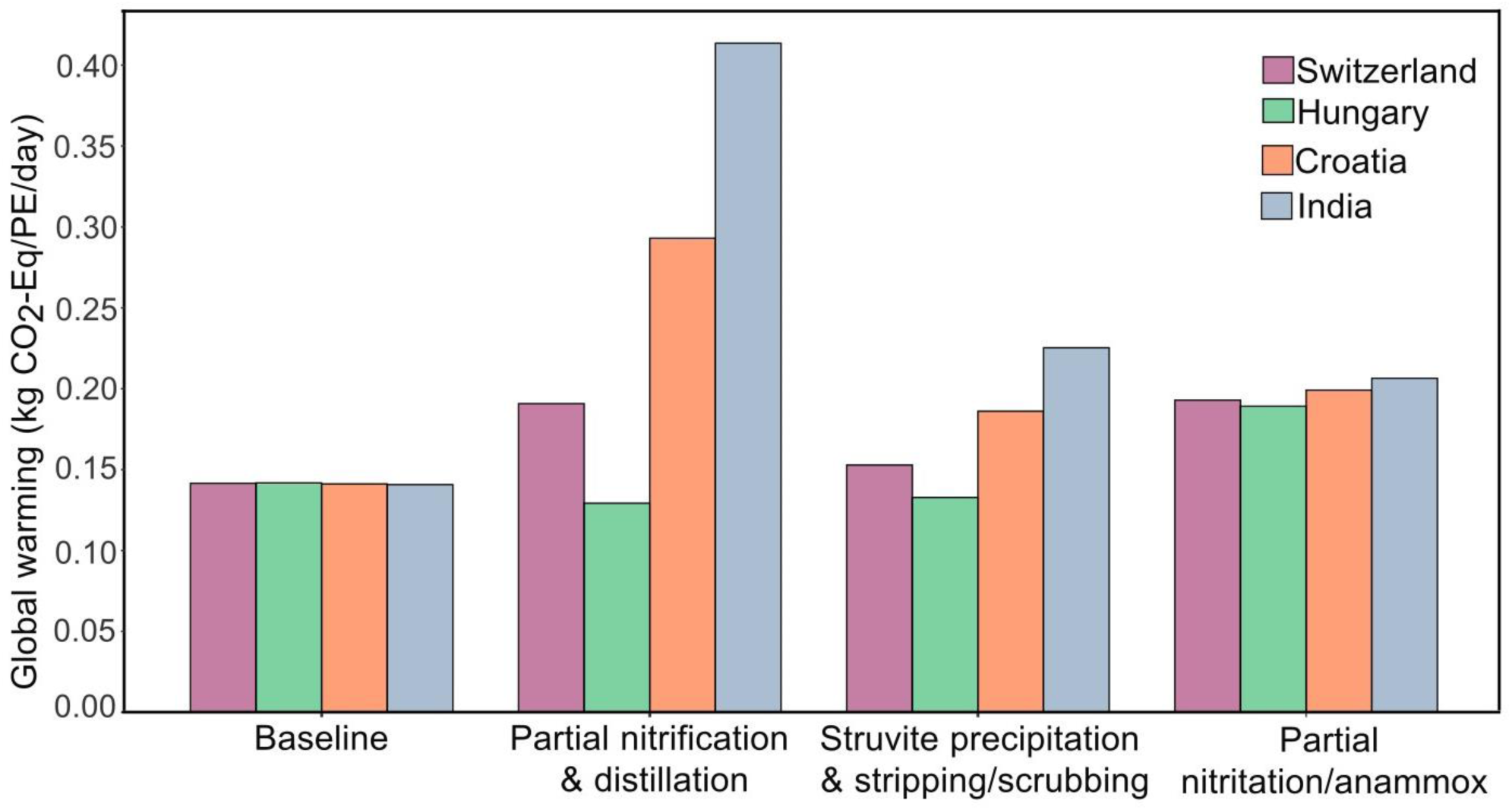
Sensitivity analysis of the studied scenarios based on the marginal electricity mixes of different countries.

At a central WWTP average of 1.1% N_2_O EF (De Haas & Andrews, 2022), both hybrid scenarios present a higher impact than the baseline at the current assumed N_2_O EF for the urine treatment and central WWTP electricity use (–0.001 kWh/PE). However, the partial nitrification & distillation and partial nitritation/anammox hybrid solutions show a lower impact at a central WWTP electricity use below 0.5 and 1 kWh/PE respectively. For the IPCC recommended EF (1.6%), partial nitrification & distillation and partial nitritation/anammox hybrid solutions presents a lower global warming than the baseline below –0.001 and 0.5 kWh/PE central WWTP electricity use respectively.

The results (Figure 8) indicate that at lower electricity carbon intensities, such as in Hungary, the partial nitrification & distillation and struvite precipitation & stripping/scrubbing hybrid scenarios have a lower global warming impact than the baseline. However, in countries with higher carbon intense electricity mix, like Croatia and India, all hybrid scenarios result in a higher impact compared to the baseline. This is due to the baseline scenario’s low electricity consumption.

The partial nitrification & distillation hybrid scenario is sensitive to different electricity mix, with its impact increasing 2 to 3 times in India compared to Switzerland and Hungary due to its high electricity use. The struvite precipitation & stripping/scrubbing scenario shows a moderate variation across countries. The partial nitritation/anammox scenario is only slightly sensitive to electricity mix variations, maintaining a less variation due to its low electricity consumption. The baseline scenario remains unaffected by changes in electricity carbon intensity, showing minimal variation across different electricity mixes.

## 4 Discussion

This study presents a comparative environmental assessment between a baseline centralized WWTP and three hybrid solutions. The findings indicate that the hybrid scenarios showed lower environmental impacts in 8 out of 10 impact categories, with the exception of global warming and marine eutrophication, showing the potential benefits of urine source-separation. Some of these benefits such as avoided fertilizer production, reduced chemicals and materials consumption in the central WWTP through urine source-separation have been reported in earlier research (Igos et al., 2017; Jimenez et al., 2015). Additionally, urine alkalinization as a pretreatment increases the environmental impact of the treatment system due to extra electricity and chemicals demand.

### 4.1 Impact of N_2_O emission on global warming

In general literature evidence provides different conclusions as to whether urine source-separation are a preferred alternative to conventional systems. While most studies conclude that urine source-separation systems have a lower global warming impact than conventional WWTPs, this study suggests that a urine source-separation is not preferred to a highly efficient central WWTP. The present research shows that one reason for these different findings (in addition to technological assumptions and modelling choices) are the emissions assumed to be occurring at the centralized WWTP. For the first time the relationship between direct N_2_O and other GHG emissions occurring at both the central WWTP and the hybrid system were quantified. Our analysis enables researchers and practitioners to plot their GHG emissions and estimate whether urine source-separation could be a lower emissions solution. With this approach this research offers a partial answer to the question at which current treatment situation a choice of source-separation would reduce GHG emission at an 80% urine source-separation.

Moreover, a combination of decentralized urine treatment with centralized WWTPs (hybrid solution) presents a promising approach to reduce the global warming impacts of conventional systems with high N₂O emissions. While the urine treatment technologies have relatively higher N₂O emissions, there is a potential for improvement due to their relatively lower technological readiness level. As studies have shown that the environmental performance of new technologies improves on maturity (Hulst et al., 2020), the Pareto frontier analysis, highlights the potential for hybrid systems to obtain a lower global warming impact than conventional WWTPs when direct N₂O emission reductions are achieved.

Among the various N₂O mitigation measures, off-gas capture and treatment from wastewater treatment systems (Duan et al., 2021) has recently gained attention. Although technologies for N₂O capture are still in early development, treatment of high-strength ammonium wastewater (urine) in sealed reactors are favorable for capturing off-gases which is uncommon for conventional WWTPs. At reduced N₂O emission levels—0.26% for partial nitrification and distillation, and 0.4% for partial nitritation/anammox—these hybrid solutions could reduce the global warming impact of 68% and 72%, respectively, of the 75 WWTPs analyzed De Haas & Andrews (2022), even for a central WWTP with a net positive energy balance. This highlights the sensitivity of urine treatment systems that use nitrification to N₂O emissions, suggesting that reducing these emissions in decentralized treatment can lower the global warming impacts of hybrid systems.

### 4.2 Influence of electricity on global warming impact

Electricity use is another factor that influences the global warming impact of WWTPs. Some studies report that electricity consumption accounts for as much as 75% of the total global warming impact in conventional WWTPs (Liao et al., 2020). However, in this study, the net positive electricity use of the central WWTP resulted in negative GHG emissions from electricity. Conventional systems with high electricity demand, such as 1.4 kWh/m³ (4 kWh/PE/day) reported by Ishii & Boyer (2015), is higher than the –0.001 kWh/PE/day of the central WWTP in this study. For such high-energy consuming systems, hybrid solutions could serve as an effective strategy to mitigate global warming impacts. The Pareto frontier analysis (Figure 7) highlights this potential. For example, the partial nitrification & distillation hybrid showed a lower global warming impact than 100% of the 75 central WWTPs (De Haas & Andrews, 2022) when central WWTP electricity use exceeded 1 kWh/PE. Similarly, the partial nitritation/anammox hybrid achieved lower global warming impacts than all conventional WWTPs studied when electricity use was above 2 kWh/PE. These findings emphasize the role of hybrid systems in reducing emissions and improving environmental performance, particularly in energy-intensive conventional WWTPs.

Additionally, the choice of electricity sources influence the environmental impact of wastewater treatment systems, particularly for energy-intensive hybrid solutions like partial nitrification & distillation and struvite precipitation & stripping/scrubbing. While this study used the Swiss electricity mix, exploring the use of cleaner, low-carbon energy sources shows the potential to reduce their global warming impact. As the world transitions toward clean energy use in the coming decades, adopting low-carbon electricity can improve the environmental performance of such systems. However, for highly efficient WWTPs, the global warming impact remains unaffected by changes in electricity sources, indicating limited benefits from clean energy for highly electricity-efficient WWTPs. This emphasizes the importance of matching energy strategies to the specific needs and efficiency levels of different treatment systems. Moreover, while optimizing electricity use in WWTPs with a decarbonized energy source may not yield significant environmental advantages, it could still provide economic benefits.

### 4.3 Opportunities for new centralized WWTPs for hybrid wastewater treatment

Urine source-separation led to higher marine eutrophication impacts in the hybrid scenarios compared to the baseline, primarily due to increased NH_4_^+^ concentration in the discharged effluent. With 80% urine separation, the COD/N ratios in the influent to the central WWTP increased, potentially allowing ordinary heterotrophic organisms to outcompete nitrifiers for COD removal. Resulting in more nitrogen being incorporated into biomass rather than supporting nitrifier growth for NH_4_^+^ oxidation. This limited NH_4_^+^ oxidation in the hybrid central WWTPs, caused higher NH_4_^+^ effluent levels. This trend aligns with findings from similar studies with 80% urine separation (Jimenez et al., 2015; Wilsenach & Van Loosdrecht, 2003).

In the partial nitrification & distillation scenario, influent nitrogen levels were already below legal limits before central WWTP treatment, suggesting that the current central WWTP model may not be the optimal choice for wastewater treatment when 80% of urine is source-separated. This creates opportunities for novel wastewater treatment plants. The current plant could be optimized for COD removal by the addition of a coagulant before primary clarification (Bisinella De Faria et al., 2015). Alternative systems such as membrane bioreactors, constructed wetland and upflow anaerobic sludge blanket can be used for high COD/N ratio wastewaters after urine separation. Future research should assess the environmental impacts and feasibility of alternative WWTP types for wastewater with high levels of source-separated urine, considering necessary infrastructure and operational modifications.

## Conclusions

This study is among the first to use a consequential LCA approach to assess the environmental impact of three decentralized urine treatment systems (partial nitrification & distillation, struvite precipitation & stripping/scrubbing, and partial nitritation/anammox) integrated with a highly efficient centralized WWTP (hybrid system) compared to a centralized WWTP (baseline) treating mixed wastewater, while also evaluating the impacts of urine alkalinization. The main conclusions of the study are:

- The hybrid systems showed improved environmental performance in 8 out of the 10 assessed impact categories, except global warming and marine eutrophication compared to the baseline. The baseline scenario’s lower global warming impact was due to the central WWTP’s low N₂O emissions and electricity consumption.
- The Pareto analysis from this study helps identify where decentralized urine treatment can reduce GHG emissions, when implemented into areas served by WWTPs with specific CO_2_-eq. emission. Thereby, helping decision makers to define the best locations for innovative urine treatment that diverts N from wastewater treatment.
- It was found that if using the N_2_O EFs of 75 WWTPs, 87% of centralized WWTPs had lower CO₂ emissions compared to partial nitrification & distillation, and 91% compared to partial nitritation/anammox hybrid solutions for a central WWTP with a net positive energy balance. However, at central WWTP electricity demands of 1 kWh/PE and 2 kWh/PE, partial nitrification & distillation and partial nitritation/anammox hybrid solutions showed a lower impact than all the studied WWTPs.
- Additionally, the analysis showed that urine alkalinization resulted in high environmental impact in 7 out of the 10 impact categories. This suggests that urine alkalinization is generally not required to lower environmental impacts of urine treatment systems, but may be relevant in systems with high NH_3_ emissions.
- The findings highlight the potential for decentralized urine treatment as a strategy to reduce global warming impacts in WWTPs with high N₂O emissions and electricity use.

## Declaration of competing interest

The authors declare that they have no known competing financial interests or personal relationships that could have appeared to influence the work reported in this paper.

## Acknowledgement

Hanson Appiah-Twum was supported by the Flemish Institute for Technological Research (VITO) and Research Foundation Flanders (FWO) (grant number 1S45425N). The authors would like to thank Kristina Bock, David de Chambrier, Nadège de Chambrier and Wenzel Gruber and Marloes Caduff for their assistance in data acquisition and contribution to this study.

## Notes

### Competing Interest Statement

The authors have declared no competing interest.

