## Supplementary material for "Environmental impact of integrating decentralized urine treatment in the urban wastewater management system: A comparative life cycle assessment"

**List of figures**

Figure S1 The layout of the defined fictitious city. The layout for the hybrid wastewater treatment and reference scenarios are on the figure's left and right, respectively. They are not drawn to scale.. 5

#### 1 Influent characteristics of the baseline and hybrid scenarios

Table S1 Influent characteristics for the studied scenarios. Influent loads are based on (Larsen & Gujer, 1996; Udert et al., 2006). Urine (80% separation) is the influent data for the decentralized urine treatment. WWTP influent concentrations for the hybrid scenarios consist of wastewater flow after urine diversion and treated urine effluent. Baseline influent characteristics are primary data of the WWTP.

| | | | | Decentralized effluent (transported to centralized WWTP) | | | | Influent to centralized WWTP | | | |
|---|---|---|---|---|---|---|---|---|---|---|---|
| Parameters | Unit | Urine | Urine (80% separation) | 20% urine to WWTP | Partial nitrification & distillation | Struvite precipitation & stripping/scrubbing | Partial nitritation/anammox | Baseline influent | Partial nitrification & distillation | Struvite precipitation & stripping/scrubbing | Partial nitritation/anammox |
| Flow rate | $m^3$/PE/day | 0.00125 | 0.0010 | 0.00025 | 0.0026 | 0.003 | 0.003 | 0.33 | 0.33 | 0.33 | 0.33 |
| COD | g/$m^3$/PE/day | 12000 | 12000 | 12000 | | 4539 | 817 | 412 | 380 | 410 | 380 |
| TN | g/$m^3$/PE/day | 9200 | 9200 | 9200 | — | 1467 | 693 | 39 | 11.2 | 22.9 | 16.7 |
| TP | g/$m^3$/PE/day | 960 | 960 | 960 | — | 5.8 | 290.5 | 5.0 | 2.0 | 2.1 | 4.4 |
| COD | g/PE/day | 15 | 12 | 3 | 2 | 12 | 2.16 | 136 | 126 | 136 | 126 |
| TN | g/PE/day | 11.5 | 9.2 | 2.30 | — | 3.9 | 1.83 | 12.9 | 3.7 | 7.6 | 5.5 |
| TP | g/PE/day | 1.2 | 0.96 | 0.24 | — | 0.02 | 0.77 | 1.6 | 0.7 | 0.7 | 1.4 |
| COD/N ratio | — | — | — | — | — | — | — | 10.5 | 33.9 | 17.9 | 22.7 |
| COD/P ratio | — | — | — | — | — | — | — | 83 | 186 | 197 | 87 |
| N/P ratio | — | — | — | — | — | — | — | 7.9 | 5.5 | 11 | 3.8 |

#### 2 Overview of the fictitious city

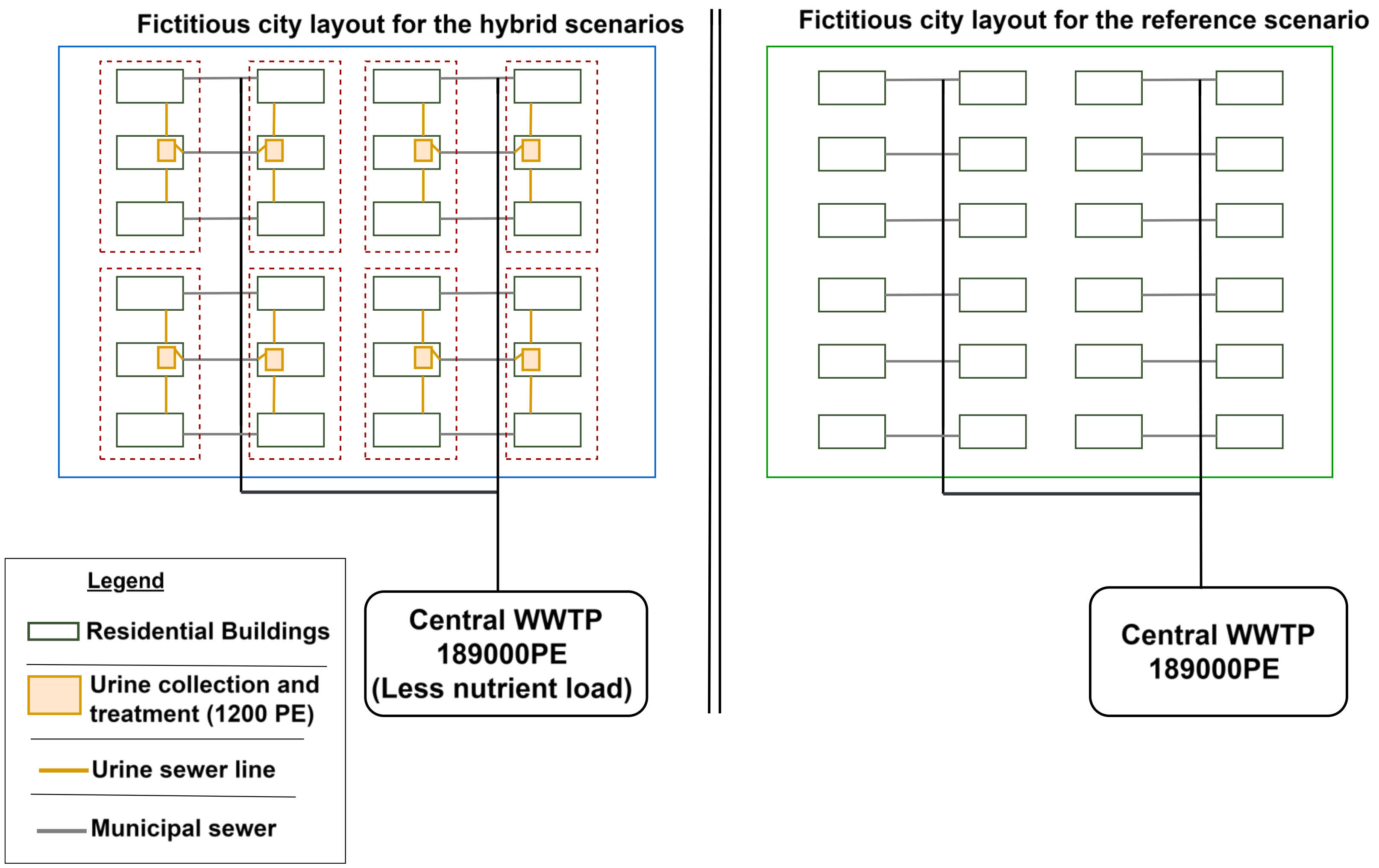

Figure S1 The layout of the defined fictitious city. The layout for the hybrid wastewater treatment and reference scenarios are on the figure's left and right, respectively. They are not drawn to scale.

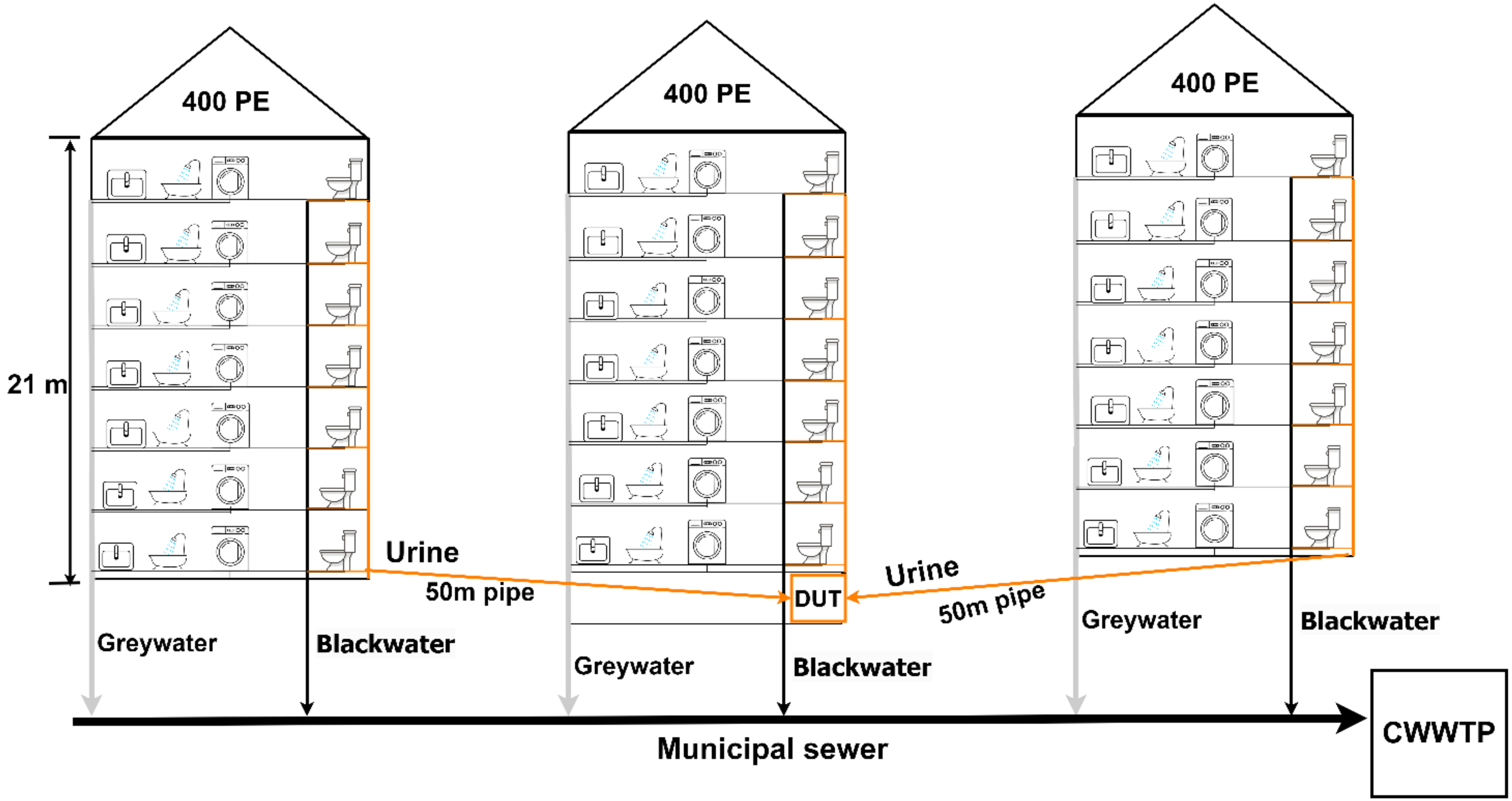

Figure S2 Building-level design in the fictitious city. DUT: decentralised urine treatment, CWWTP: centralised wastewater treatment plant.

### 3 Plumbing estimation for urine transportation

#### 3.1 Methodological approach for the piping estimation

The piping infrastructure for urine transportation was modelled using the International Plumbing Code (IPC) (International Plumbing Code, 2024). We used the assumed characteristics of the theoretical city to design the layout of the plumbing infrastructure; 7-storey buildings of height 21m, accommodating 400 inhabitants and an average of 2.2 inhabitants per dwelling corresponding to 182 dwellings. Each floor has a height of 3m and there are 26 apartments on each floor. It is assumed that each apartment has one source-separating toilet and there is an assumed distance of 5m from each toilet. The plumbing plan consisting of fixtures on each floor, horizontal fixture branches, vertical stack pipes and offsets were drawn (Figure S3). Venting pipes were excluded. We then determined the drainage fixture unit (DFU) for urine only which is 2, based on the IPC. DFU measures the drainage capacity of a plumbing fixture into the drainage system. The total DFU for the horizontal branch pipe from all seven floors were calculated. The diameter of each horizontal fixture branch pipe was determined (Table S2) using the IPC reference tables. The peak flow hour was taken into consideration. The DFU for all the horizontal pipes in the building was calculated and then referred to the IPC to determine the diameter of the vertical stack pipe. We then determined the offset pipes' size and the vertical stack's length. We assumed that the pipes were schedule 40 type based on ANSI pipe schedule chart (Yaro, 2024) to estimate the weight of the pipes. We assumed a 50 year lifetime for the pipes. The total pipe estimation is shown in **Error! Reference source not found.**.

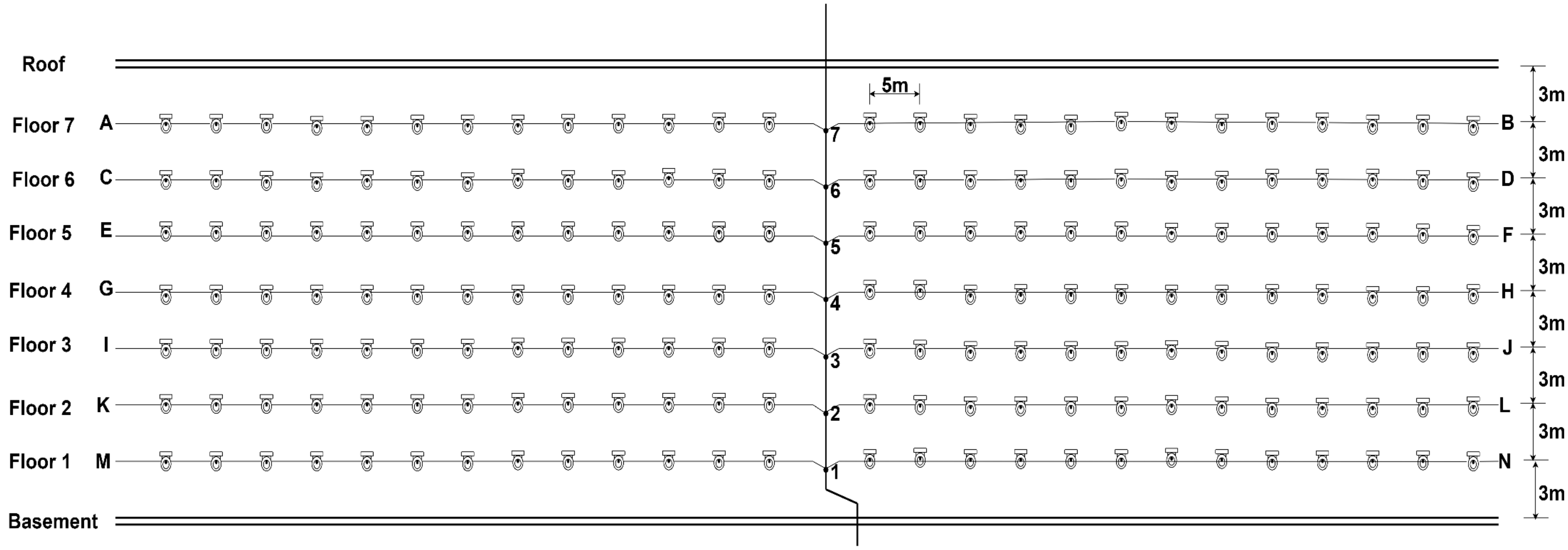

Figure S3 Layout of the plumbing design for urine transportation

##### 3.2 Total fixture unit of the horizontal branch of all floors

Table S2 Total fixture unit of the horizontal branch of all floors

| Branch | Floor | Fixture unit (DFU) | Quantity of fixtures | Total DFU | Total DFU of branch | Horizontal fixture Length (m) | Pipe diameter in/ (mm) |
|---|---|---|---|---|---|---|---|
| A | 7 | 2 | 13 | 26 | | 65 | 100 |
| B | | 2 | 13 | 26 | 52 | 65 | 100 |
| C | 6 | 2 | 13 | 26 | | 65 | 100 |
| D | | 2 | 13 | 26 | 52 | 65 | 100 |
| E | 5 | 2 | 13 | 26 | | 65 | 100 |
| F | | 2 | 13 | 26 | 52 | 65 | 100 |
| G | 4 | 2 | 13 | 26 | | 65 | 100 |
| H | | 2 | 13 | 26 | 52 | 65 | 100 |
| I | 3 | 2 | 13 | 26 | | 65 | 100 |
| J | | 2 | 13 | 26 | 52 | 65 | 100 |
| K | 2 | 2 | 13 | 26 | | 65 | 100 |
| L | | 2 | 13 | 26 | 52 | 65 | 100 |
| M | 1 | 2 | 13 | 26 | | 65 | 100 |
| N | | 2 | 13 | 26 | 52 | 65 | 100 |
| | | | 182 | Total | **364** | **910** | |

### 4 Results

#### 4.1 Total pipes needed for urine transportation

Table S3 Estimated total pipes for urine transportation

| | Horizontal fixture branch | Vertical stack | Offset | Total kg/ 400 PE | Total kg/ PE/day |
|---|---|---|---|---|---|
| Diameter (mm) | 100 | 100 | 125 | — | — |
| Length (m) | 910 | 23 | 2 | — | — |
| Weight (kg/m) | 16.07 | 16.07 | 11.57 | — | — |
| Total weight | 14623.7 | 369.61 | 23.14 | 15016.45 | 0.00206 |

The piping infrastructure was modelled for 400 PE and normalized to the functional unit of PE/day assuming a 50-year lifetime for the pipes. PVC material was used for the in-building pipes.

Additionally, we assumed a 50m underground residential sewer line for urine transport between the three buildings for treatment. Since the treatment is in the basement of one of the three buildings, we will need two of the 50m lines for this purpose, which is equivalent to a 100m sewer length. We assumed a 50-year lifetime for the residential sewer line. The Ecoinvent process, "residential sewer grid construction, 0.087 km, CH" was used for the underground residential sewer line for urine transport between the three buildings for treatment.

#### 4.2 Life cycle inventory of the decentralized urine treatment systems' infrastructure

Table S4 A detailed inventory of the infrastructural demand for the decentralized urine treatment systems

| Unit processes | Material composition | Partial nitrification & distillation (without alkalinization) | Partial nitrification & distillation [$Ca(OH)_2$ alkalinization] | Partial nitrification & distillation (Electro-chemical alkalinization) | Struvite precipitation & stripping/ scrubbing | Partial nitritation/ anammox | Unit (PE/day) | Comment |
|---|---|---|---|---|---|---|---|---|
| Collection tank | HDPE | 0.00013 | 0.00013 | 0.00013 | 0.00013 | 0.00013 | kg | 20 $m^3$. This is equivalent to 1145 kg. Material made of HDPE. Lifetime of 20 years |
| Pre-stabilisation | Steel | — | — | 4.52E-05 | — | — | kg | Assume the skid without the collection tank has a mass of 400 kg of which 99% is stainless steel and 1% of PVC. 20 year life assumed |
| | PVC | — | — | 4.57E-07 | — | — | kg | |
| Nitrification reactor | HDPE | 3.8E-05 | 3.8E-05 | 3.8E-05 | — | 3.8E-05 | kg | 3 tanks of which are Deshoust PE 4000 D type. Weighs 235 kg each. The PE weighs 111kg while the steel weighs 124kg and has a volume of 4000 L. Lifetime of 20 years |
| | Steel | 4.18E-05 | 4.18E-05 | 4.18E-05 | — | 4.18E-05 | kg | Galvanized steel braces. Made of steel and coated with zinc. Area of coating estimated |
| | Zinc coating | 2.67E-06 | 2.67E-06 | 2.67E-06 | — | 2.67E-06 | m2 | |
| Granular activated carbon | Composite fibreglass | 5.14E-05 | 5.14E-05 | 5.14E-05 | — | 5.14E-05 | kg | Lifetime of 20 years |
| Distiller | Steel | 0.00011 | 0.00011 | 0.00011 | — | — | kg | Prowadest 120 model with a weight of 960 kg which is 99% super duplex steel ( has lower amount of Ni and Mo). Lifetime of 20 years based on personal communication |
| | PVC | 1.1E-06 | 1.1E-06 | 1.1E-06 | — | — | kg | All forms of tubings or plastics in the set-up |

| | | | | | | | | |
|---|---|---|---|---|---|---|---|---|
| Intermediate storage tank | HDPE | 1.23E-05 | 1.23E-05 | 1.23E-05 | — | — | kg | 2 tanks which are Deshoust PE 2500 D type for this FU. Weighs 115 kg each. The PE weighs 47% while the steel weighs 53% of the total weight and has a volume of 4000 L. Lifetime of 20 years |
| | Steel | 1.37E-05 | 1.37E-05 | 1.37E-05 | — | — | kg | Galvanized steel braces. Same comment as in the Nitrification reactor |
| | Zinc | 8.76E-07 | 8.76E-07 | 8.76E-07 | — | — | m2 | |
| Struvite reactor | Composite fiber glass | — | — | — | 5.14E-05 | — | kg | Lifetime of 20 years |
| Stripper/scrubber | Polypropylen e | — | — | — | 1.85E-06 | — | kg | Lifetime of 20 years |
| | Steel | — | — | — | 4.12E-05 | — | kg | Lifetime of 20 years |
| | PVC | — | — | — | 0.00039 | — | kg | Lifetime of 20 years |
| Nitric acid storage | Polyethylene | — | — | — | 0.00014 | — | kg | Lifetime of 20 years |

#### 4.3 Life cycle inventory of the central wastewater treatment plants

Table S5 Life cycle inventory of the central wastewater treatment plants of the studied scenarios

| Treatment configuration | Flow description | Baseline WWTP | Partial nitrification & distillation (central WWTP) | Struvite precipitation & stripping/scrubbing (central WWTP) | Partial nitritation /anammox (central WWTP) | Unit | Source/ Comment | Ecoinvent Database process/flow used |
|---|---|---|---|---|---|---|---|---|
| Wastewater treatment | Electricity (primary treatment) | 0.0026 | 0.0026 | 0.0026 | 0.0026 | kWh | Primary data | market for electricity, medium, CH |
| | $FeCl_3$ dosage (P removal) | 0.0012 | — | — | — | kgFe | Primary data | market for iron(III) chloride, without water, in 40% solution state |
| | Electricity (biology treatment) | 0.0108 | 0.0108 | 0.0108 | 0.0108 | kWh | Primary data | market for electricity, medium, CH |
| | Electricity (aeration) | 0.0194 | 0.0103 | 0.0131 | 0.0120 | kWh | Simulation data for the hybrid WWTPs | market for electricity, medium, CH |
| | $N_2O$ emission (biology treatment) | 0.3 | 0.3 | 0.3 | 0.3 | % influent TN | Primary data | Dinitrogen monoxide ('air') |
| | $CH_4$ emission (biology treatment) | 0.1 | 0.1 | 0.1 | 0.1 | % influent COD | Wenzel, 2021 | methane, non fossil ('air') |
| | Activated carbon | 0.00230 | 0.00112 | 0.00112 | 0.00112 | kg | Hybrid WWTPs have been linearly scaled with a 49% reduction factor with the motivation that, 64% of active micropollutants in wastewater are from urine (Lienert et al., 2007) | market for activated carbon, granular |
| | $FeCl_3$ dosage (Micropollutant removal) | 0.00025 | 0.00012 | 0.00012 | 0.00012 | kgFe | | market for iron(III) chloride, without water, in 40% solution state |
| | Polymer (micropollutant removal) | 0.00065 | 0.00032 | 0.00032 | 0.00032 | kg | | market for polyacrylamide, GLO |
| | Electricity (micropollutant removal) | 0.0063 | 0.00305 | 0.00305 | 0.00305 | kWh | | market for electricity, medium, CH |
| | Electricity (filtration) | 0.0041 | 0.00406 | 0.00406 | 0.00406 | kWh | Primary data | market for electricity, medium, CH |

| | | | | | | | | |
|---|---|---|---|---|---|---|---|---|
| **Sludge management** | Electricity (anaerobic digestion) | 0.0099 | 0.0099 | 0.0099 | 0.0099 | kWh | Primary data | market for electricity, medium, CH |
| | $CH_4$ emission | 0.9 | 0.9 | 0.9 | 0.9 | % influent COD | (Gruber, 2021) | methane, non fossil ('air') |
| | Heat consumption (CHP) | 0.0367 | 0.0367 | 0.0367 | 0.0367 | — | Primary data | market for heat, district or industrial, natural gas, CH |
| | Electricity production from CHP | 0.0597 | 0.0597 | 0.0597 | 0.0597 | kWh | Primary data. Avoided electricity production | market for electricity, medium, CH |
| | Heat production from CHP | 0.0438 | 0.0438 | 0.0438 | 0.0438 | kWh | Primary data. Avoided heat production | market for heat, district or industrial, natural gas, CH |
| **Sludge reject water treatment** | Electricity (reject water treatment) | 0.0011 | 0.0011 | 0.0011 | 0.0011 | kWh | Primary data | market for electricity, medium, CH |
| | $N_2O$ emission (reject water treatment) | 1.8 | 1.8 | 1.8 | 1.8 | % N in reject water | (Dieziger et al., 2023) | Dinitrogen monoxide ('air') |
| **Facility management** | Electricity | 0.0047 | 0.0047 | 0.0047 | 0.0047 | kWh | Primary data | market for electricity, medium, CH |
| | Heat | 0.0070 | 0.0070 | 0.0070 | 0.0070 | kWh | Primary data | market for heat, district or industrial, natural gas, CH |
| **External sludge and screening management** | Distance to incineration | 20 | 20 | 20 | 20 | km | Personal communication | market for transport, freight, lorry 16-32 metric ton, EURO4, RER |
| | Distance to landfill | 13 | 13 | 13 | 13 | km | Personal communication | market for transport, freight, lorry 16-32 metric ton, EURO4, RER |
| | Screenings incineration | 0.0061 | 0.0061 | 0.0061 | 0.0061 | kg | Primary data | Treatment of municipal solid waste, incineration, CH |
| | Sludge incineration | 0.0183 | 0.0183 | 0.0183 | 0.0183 | kg | Database process was modified to include the $N_2O$ emissions | treatment of raw sewage sludge, municipal incineration, CH |
| | $N_2O$ emissions (sludge incineration) | 3.8 | 3.8 | 3.8 | 3.8 | % N in sludge | (Gruber, 2021) | Dinitrogen monoxide ('air') |

| | | | | | | | | |
|---|---|---|---|---|---|---|---|---|
| | Landfilling (incinerated waste and sand) | 0.0029 | 0.0029 | 0.0029 | 0.0029 | kg | 90% mass reduction is assumed for incinerated sludge and screenings. Mass of sand is based on primary data | treatment of inert waste, sanitary landfill, CH |

#### 4.4 Operational life cycle inventory of the partial nitrification & distillation decentralized urine treatment scenarios including the alkalinization scenarios

Table S6 Operational life cycle inventory of the partial nitrification and distillation decentralized urine treatment scenarios including the pre-stabilisation scenarios

| Unit process | Flow description | [a] Without akalinization | $Ca(OH)_2$ alkalinization | Electrochemical alkalinization | Unit | Source/ Comment | Ecoinvent Database process/flow used |
|---|---|---|---|---|---|---|---|
| **Collection tank** | $NH_3$ (emission) | 0.50 | — | — | % influent N | $NH_3$ emissions only occur in the scenario without alkalinization. The alkalinization scenarios have no $NH_3$ emissions. | Ammonia ('air') |
| | $P_2O_5$ fertilizer | 20.00 | 100.00t | 70.00 | % influent P | Spontaneous P precipitation in scenario without alkalinization (Udert et al., 2003). \| P is completely removed in $Ca(OH)_2$ alkalinization (Randall et al., 2016) \| 70% P is removed in electrochemical alkalinization. | market for single superphosphate, RER |
| | Citric acid (10%) | 40.00 | 40.00 | — | ml (bi-weekly) | 40 ml of 10% citric acid for bi-weekly toilet cleaning based on personal communication. In electrochemical alkalinization, citric acid was not considered because the electrolyte can be used for this purpose. | Market for citric acid, GLO |
| | Electricity | — | 9.68E-06 | 0.037 | kWh | Electricity demand for electrochemical alkalinization calculated from Mufunde and Randall, 2022 (Mufunde & Randall, 2022). Urine is stirred at 2L intervals at 500 rpm for 15 s.\| electrochemical alkalinization - 14 Wh/L urine (De Paepe et al., 2020) | market for electricity, medium voltage,CH |
| **Nitrification reactor** | $N_2O$ (emission) | 0.70 | 0.70 | 0.70 | % influent N | (Faust et al., 2022) | Dinitrogen monoxide ('air') |
| | Electricity (Aeration) | 0.0156 | 0.0267 | 0.0187 | kWh | 5.9 Wh/L urine (Faust et al., 2022) \| $Ca(OH)_2$ and electrochemical alkalinization estimated based 86% and 60% nitrification respectively | market for electricity, medium voltage,CH |
| **Intermediate storage tank** | $N_2O$ (emission) | 0.80 | 0.80 | 0.80 | % influent N | (Faust et al., 2022) | Dinitrogen monoxide ('air',) |

| | | | | | | | |
|---|---|---|---|---|---|---|---|
| **AC reactor** | Activated carbon | 0.5690 | 0.5690 | 0.5690 | g/L urine | (Köpping et al., 2020) | market for activated carbon, granular,GLO |
| **Distillation** | Electricity | 0.211 | 0.211 | 0.211 | kWh | 80kWh/m$^3$ urine (Personal communication) | market for electricity, medium voltage,CH |
| | N fertilizer recovered | 98 | 98 | 98 | % | — | market for ammonium nitrate, RER |
| | P recovered | 80 | — | 30 | % | Remaining P after precipitation in the collection tank | market for single superphosphate, RER |
| **Pumping** | Electricity | 0.0032 | 0.0032 | 0.0032 | kWh/L urine | (Faust et al., 2022) | market for electricity, medium voltage,CH |
| **Process control** | Electricity | 0.0032 | 0.0032 | 0.0032 | kWh/L urine | (Faust et al., 2022) | market for electricity, medium voltage,CH |
| **Fertilizer transport to field** | | 20.00 | 20.00 | 20.00 | km | Monthly transportation at 20km assumed | market for transport, freight, lorry 28 metric ton, fatty acid methyl ester 100%, CH |

[a] The inventory for the scenario without alkalinization is also used in the decentralised urine treatment for the partial nitrification & distillation hybrid wastewater treatment.

#### 4.5 Operational life cycle inventory of the struvite precipitation & stripping/scrubbing decentralized urine treatment scenario

Table S7 Operational life cycle inventory of the struvite precipitation & stripping scrubbing decentralized urine treatment scenario

| Unit process | Flow description | Amount | Unit | Source/ Comment | Ecoinvent Database process/flow used |
|---|---|---|---|---|---|
| **Collection tank** | $NH_3$ (emissions) | 0.5 | % | — | Ammonia ('air') |
| | P recovered | 20 | % influent P | Spontaneous P precipitation (Udert et al 2003b). | market for single superphosphate, RER |
| | Citric acid (10%) | 40 | ml (bi-weekly) | 40 ml of 10% citric acid for bi-weekly toilet cleaning based on personal communication. In Nit-Dist+ES, citric acid was not considered because the electrolyte can be used for this purpose. | Market for citric acid, GLO |
| **Struvite precipitation reactor** | Electricity for precipitation | 70 | kWh/kg P as struvite | (Antonini et al., 2011) | market for electricity, medium voltage, CH |
| | Mg: P ratio | 1.5:1 | | (Antonini et al., 2011) | market for magnesium oxide, GLO |
| | P recovered in struvite | 98 | % P in influent flow to struvite reactor | (Antonini et al., 2011) | market for single superphosphate, RER |
| | Electricity for struvite drying | 0.783 | kWh/kg struvite | (Hilton et al., 2021) | market for electricity, medium voltage, CH |
| **Stripping/ scrubber unit** | $HNO_3$:N | 2:1 | | Personal communication | market for nitric acid, without water, in 50% solution state, RER w/o RU |
| | Electricity | 5.701 | kWh/kgN stripped | Personal communication | market for electricity, medium voltage,CH |
| | $NH_3$ (emissions) | 0.000031 | % | $NH_3$ emission is negligible, Personal communication. | Ammonia ( emissions to air) |
| | N fertilizer recovered | 54 | % N influent | 56% N removal efficiency with an Ammonium nitrate product (7.7% N). Personal communication | market for ammonium nitrate, RER |
| **Activated carbon reactor** | Activated carbon | 0.5690 | g/L urine | (Köpping et al., 2020) | market for activated carbon, granular,GLO |
| **Fertilizer transport to field** | | 20.00 | km | Monthly transportation at 20km assumed | market for transport, freight, lorry 28 metric ton, fatty acid methyl ester 100%, CH |

#### 4.6 Operational life cycle inventory of the partial nitritation/anammox decentralized urine treatment scenarios

Table S8 Operational life cycle inventory of the partial nitritation/anammox decentralized urine treatment scenarios

| Unit process | Flow description | Amount | Unit | Source/ Comment | Ecoinvent Database process/flow used |
|---|---|---|---|---|---|
| **Collection tank** | $NH_3$ (emission) | 0.50 | % influent N | — | Ammonia ('air') |
| | P recovered | 20.00 | % influent P | Spontaneous P precipitation (Udert et al 2003b). | market for single superphosphate, RER |
| | Citric acid | 40.00 | ml (bi-weekly) | 40 ml of 10% citric acid for bi-weekly toilet cleaning based on personal communication. In Nit-Dist+ES, citric acid was not considered because the electrolyte can be used for this purpose. | Market for citric acid, GLO |
| **Partial nitritation/anammox** | Electricity | 1.156 | kWh/kg N | Estimated from Faust et al, 2022 with 60% nitrification | market for electricity, medium voltage,CH |
| | $N_2O$ | 1.8% | % influent N | (Dieziger et al., 2023) | Dinitrogen monoxide ('air') |
| **AC reactor** | Activated carbon | 0.5690 | g/L urine | (Köpping et al., 2020) | market for activated carbon, granular,GLO |
| **Pumping** | Electricity | 0.0032 | kWh/L urine | (Faust et al., 2022) | market for electricity, medium voltage,CH |
| **Process control** | Electricity | 0.0032 | kWh/L urine | (Faust et al., 2022) | market for electricity, medium voltage,CH |
| **Fertilizer transport to the field** | | 20.00 | km | Monthly transportation at 20km assumed | market for transport, freight, lorry 28 metric ton, fatty acid methyl ester 100%, CH |

#### 4.7 Modified model calibration parameters

Table S9 Details of modified model calibration parameters

| Configuration | Flow description | Baseline WWTP | Partial nitrification & distillation (central WWTP) | Struvite precipitation & stripping/scrubbing (central WWTP) | Partial nitritation/anammox (central WWTP)) | Unit |
|---|---|---|---|---|---|---|
| **General conditions** | Temperature | 20 | 20 | 20 | 20 | °C |
| | pH | 7.2 | 7.2 | 7.2 | 7.2 | pHunit |
| **Primary clarifier** | Total suspended solids percent removal | 50 | 50 | 50 | 50 | % |
| | Sludge flow | 171 | 171 | 171 | 171 | m3/d |
| **Anaerobic reactor** | Volume per train | 7950 | 7950 | 7950 | 7950 | m3 |
| | Air flow @ standard conditions (NTP: 20 °C, 1 atm) | 0 | 0 | 0 | 0 | m3/d at NTP |
| **Anoxic reactor** | Volume per train | 3900 | 3900 | 3900 | 3900 | m3 |
| | Air flow @ standard conditions (NTP: 20 °C, 1 atm) | 0 | 0 | 0 | 0 | m3/d at NTP |
| **Aerobic reactor 1** | Volume per train | 6475 | 6475 | 6475 | 6475 | m3 |
| | Air flow @ standard conditions (NTP: 20 °C, 1 atm) | 95462.6 | 60000 | 62562.4 | 68000 | m3/d at NTP |
| | Alpha (wastewater/clean water) factor | 0.8 | 0.8 | 0.8 | 0.8 | — |
| | Diffuser fouling factor | 0.9 | 0.9 | 0.9 | 0.9 | — |
| | Diffuser floor density (diffuser area/tank area) | 0.21 | 0.21 | 0.21 | 0.21 | m2/m2 |
| | Area per diffuser | 0.065 | 0.065 | 0.065 | 0.065 | m2 |
| | Intercept in SSOTE correlation | 8 | 8 | 8 | 8 | %/m |
| | Asymptote in SSOTE correlation | 7 | 7 | 7 | 7 | %/m |
| | Blower efficiency | 68 | 68 | 68 | 68 | % |
| **Aerobic reactor 2** | Volume per train | 5410 | 5410 | 5410 | 5410 | m3 |
| | Air flow @ standard conditions (NTP: 20 °C, 1 atm) | 61005.7 | 20000 | 26000 | 28500 | m3/d at NTP |
| | Alpha (wastewater/clean water) factor | 0.8 | 0.8 | 0.8 | 0.8 | — |
| | Diffuser fouling factor | 0.9 | 0.9 | 0.9 | 0.9 | — |
| | Diffuser floor density (diffuser area/tank area) | 0.19 | 0.19 | 0.19 | 0.19 | m2/m2 |
| | Area per diffuser | 0.065 | 0.065 | 0.065 | 0.065 | m2 |
| | Intercept in SSOTE correlation | 8 | 8 | 8 | 8 | %/m |

| | | | | | | |
|---|---|---|---|---|---|---|
| | Asymptote in SSOTE correlation | 7 | 7 | 7 | 7 | %/m |
| | Blower efficiency | 68 | 68 | 68 | 68 | % |
| **Aerobic reactor 3** | Volume per train | 5410 | 5410 | 5410 | 5410 | m3 |
| | Air flow @ standard conditions (NTP: 20 °C, 1 atm) | 18073 | 10000 | 10000 | 10000 | m3/d at NTP |
| | Alpha (wastewater/clean water) factor | 0.8 | 0.8 | 0.8 | 0.8 | — |
| | Diffuser fouling factor | 0.9 | 0.9 | 0.9 | 0.9 | — |
| | Diffuser floor density (diffuser area/tank area) | 0.19 | 0.19 | 0.19 | 0.19 | m2/m2 |
| | Area per diffuser | 0.065 | 0.065 | 0.065 | 0.065 | m2 |
| | Intercept in SSOTE correlation | 8 | 8 | 8 | 8 | %/m |
| | Asymptote in SSOTE correlation | 7 | 7 | 7 | 7 | %/m |
| | Blower efficiency | 68 | 68 | 68 | 68 | % |
| **Aerobic reactor 4** | Volume per train | 2415 | 2415 | 2415 | 2415 | m3 |
| | Air flow @ standard conditions (NTP: 20 °C, 1 atm) | 4040 | 5000 | 5000 | 5000 | m3/d at NTP |
| | Alpha (wastewater/clean water) factor | 0.8 | 0.8 | 0.8 | 0.8 | — |
| | Diffuser fouling factor | 0.9 | 0.9 | 0.9 | 0.9 | — |
| | Diffuser floor density (diffuser area/tank area) | 0.17 | 0.17 | 0.17 | 0.17 | m2/m2 |
| | Area per diffuser | 0.065 | 0.065 | 0.065 | 0.065 | m2 |
| | Intercept in SSOTE correlation | 8 | 8 | 8 | 8 | %/m |
| | Asymptote in SSOTE correlation | 7 | 7 | 7 | 7 | %/m |
| | Blower efficiency | 68 | 68 | 68 | 68 | % |
| **Pumped flows** | Internal recycling (IR) | 47526 | 47526 | 47526 | 47526 | m3/d |
| | Return activated sludge (RAS) | 47073.3 | 47073.3 | 47073.3 | 47073.3 | m3/d |
| | Waste activated sludge (WAS) | 1133.7 | 1133.7 | 1133.7 | 1133.7 | m3/d |
| **Iron dosage** | Fe mass flow | 220 | — | — | — | kg Fe/d |
| | Iron concentration | 190000 | — | 190000 | 190000 | g Fe/m3 |
| | Average velocity gradient at dosage point | 100 | — | 100 | 100 | 1/s |
| **Secondary clarifier** | Total volume per train | 13500 | 13500 | 13500 | 13500 | m3 |
| | Sludge flow | 48207 | 48207 | 48207 | 48207 | m3/d |
| | Effluent solids | 1 | 1 | 1 | 1 | g/m3 |

#### 4.8 Electricity balance for the urine alkalinization scenarios

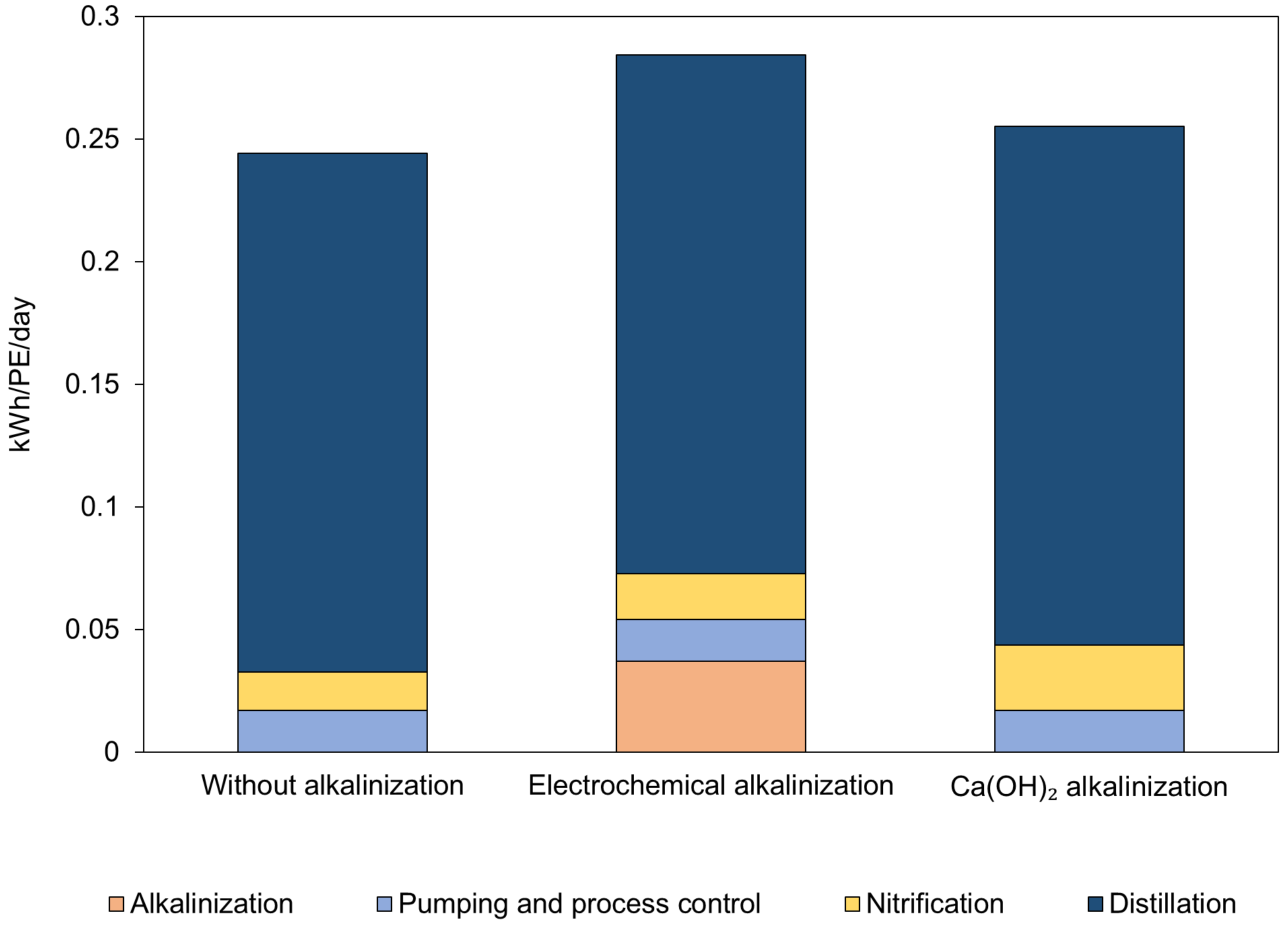

Figure S4 Energy balance of the urine alkalinization scenarios

#### 4.9 Average concentration of nitrifiers

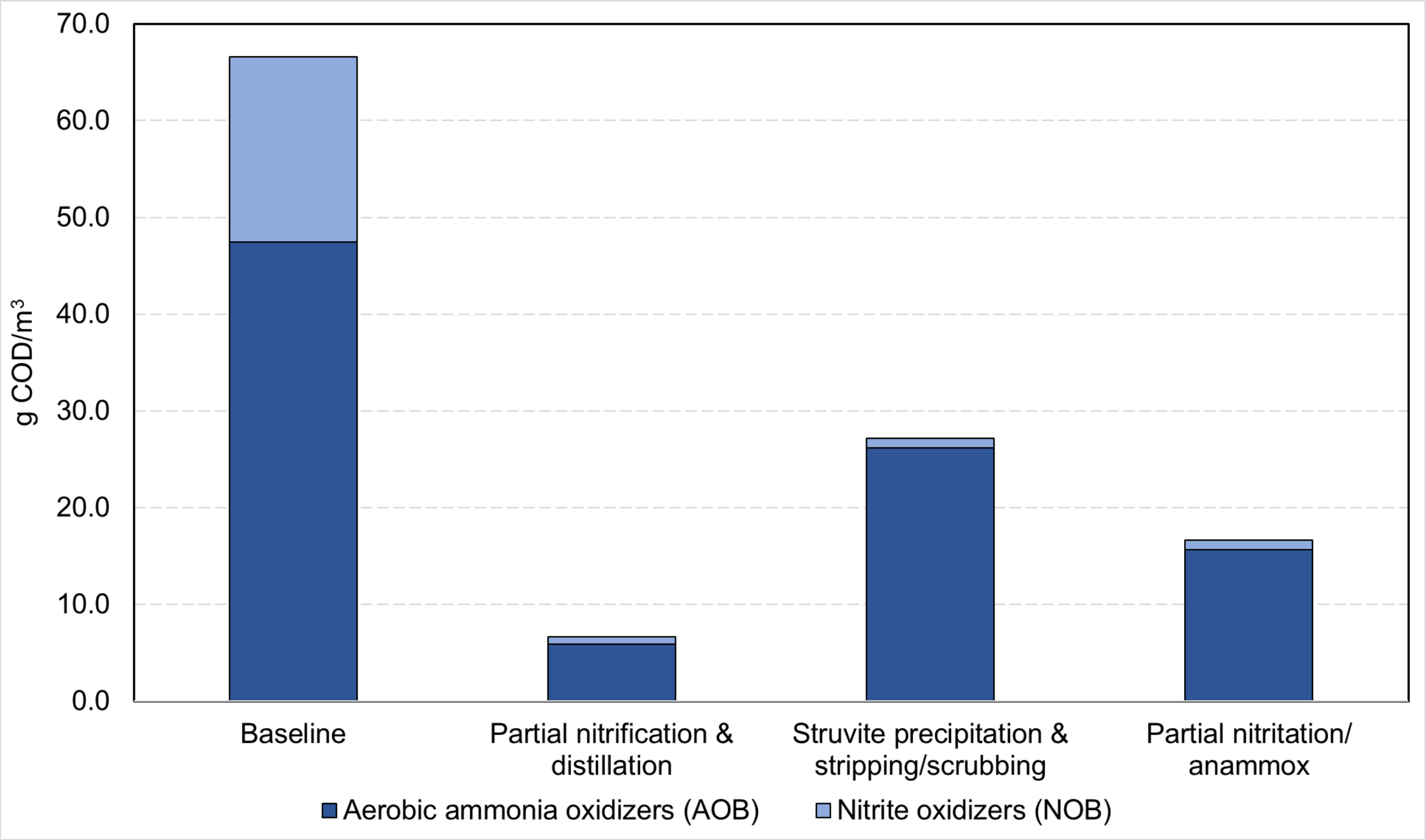

Figure S5 Average concentration of nitrifiers in the central WWTP across the scenarios

##### 4.10 Impact assessment results of the baseline and hybrid wastewater treatment scenarios

Table S10 Impact assessment results of the baseline and hybrid scenarios

| Impact categories | Unit | Baseline | Partial nitrification & distillation | Struvite precipitation & stripping/scrubbing | Partial nitritation/anammox |
|---|---|---|---|---|---|
| Acidification (Terrestrial) | kg $SO_2$-eq/ PE/day | 8.35E-05 | 5.32E-05 | 2.88E-04 | 1.89E-04 |
| Global warming | kg $CO_2$-eq/PE/day | 0.142 | 0.191 | 0.153 | 0.193 |
| Ecotoxicity (freshwater) | kg 1,4-DCB-eq/PE/day | 0.006 | 0.003 | 0.004 | 0.003 |
| Ecotoxicity (marine) | kg 1,4-DCB-eq/PE/day | 0.008 | 0.004 | 0.005 | 0.004 |
| Fossil depletion | kg oil-eq/PE/day | 0.011 | 0.014 | 0.018 | 0.010 |
| Eutrophication (freshwater) | kg P-eq/PE/day | 8.38E-05 | 3.62E-05 | 4.08E-05 | 4.56E-05 |
| Eutrophication (marine) | kg N-eq/PE/day | 1.31E-04 | 1.42E-04 | 1.38E-04 | 1.43E-04 |
| Minerals depletion | kg Cu-eq/PE/day | 6.96E-04 | -1.82E-04 | 2.90E-03 | 3.02E-05 |
| Ozone depletion | kg CFC11-eq/PE/day | 2.32E-06 | 3.49E-06 | 1.79E-06 | 4.40E-06 |
| Particulate matter formation | kg PM2.5-eq/PE/day | 3.79E-05 | 2.02E-05 | 5.91E-05 | 4.88E-05 |

The results of the hybrid treatment scenarios include the operational impact of decentralized urine treatment and central WWTP, as well as the impact of the urine treatment infrastructure and piping infrastructure for urine transportation.

#### 4.11 Impact assessment results of the urine alkalinization scenarios

Table S11 Impact assessment results of the urine alkalinization scenarios

| Impact categories | **Unit** | **Without alkalinization** | **Electrochemical alkalinization** | **Calcium hydroxide alkalinization** |
|---|---|---|---|---|
| Acidification (Terrestrial) | kg $SO_2$-eq/ PE/day | 8.06E-06 | -9.91E-05 | -9.15E-05 |
| Global warming | kg $CO_2$-eq/PE/day | 1.05E-01 | 1.17E-01 | 1.32E-01 |
| Ecotoxicity (freshwater) | kg 1,4-DCB-eq/PE/day | 9.00E-04 | 1.36E-03 | 9.93E-04 |
| Ecotoxicity (marine) | kg 1,4-DCB-eq/PE/day | 9.26E-04 | 1.50E-03 | 1.04E-03 |
| Fossil depletion | kg oil-eq/PE/day | 1.19E-02 | 1.48E-02 | 1.51E-02 |
| Eutrophication (freshwater) | kg P-eq/PE/day | -6.87E-06 | -6.91E-06 | -8.08E-06 |
| Eutrophication (marine) | kg N-eq/PE/day | -2.75E-06 | -3.05E-06 | -2.73E-06 |
| Metals depletion | kg Cu-eq/PE/day | -2.36E-04 | -1.66E-04 | -2.22E-04 |
| Ozone depletion | kg CFC11-eq/PE/day | 2.25E-06 | 2.31E-06 | 2.26E-06 |
| Particulate matter formation | kg PM2.5-eq/PE/day | 3.46E-06 | -9.37E-06 | -1.01E-05 |
